# Lipophilic Nanocrystal Prodrug-Release Defines the Extended Pharmacokinetic Profiles of a Year-Long Cabotegravir

**DOI:** 10.1101/2021.01.18.427147

**Authors:** Nagsen Gautam, JoEllyn M. McMillan, Devendra Kumar, Aditya N. Bade, Qiaoyu Pan, Tanmay A. Kulkarni, Wenkuan Li, Nathan A Smith, Bhagya L. Dyavar Shetty, Brady Sillman, Adam Szlachetka, Benson J. Edagwa, Howard E. Gendelman, Yazen Alnouti

**Affiliations:** Department of Pharmaceutical Sciences, University of Nebraska Medical Center, Omaha, NE 68198 USA; Department of Pharmacology and Experimental Neuroscience, University of Nebraska Medical Center, Omaha, NE 68198 USA; Nebraska Nanomedicine Production Plant, University of Nebraska Medical Center, Omaha, NE 68198 USA

**Keywords:** cabotegravir, prodrug, long acting slow effective release antiretroviral therapy, nanoformulations, pharmacokinetics, macrophage depots, toxicology

## Abstract

A single, once every eight-week cabotegravir (CAB) long-acting parenteral is more effective than daily oral emtricitabine and tenofovir disoproxil fumarate in preventing human immunodeficiency virus type one (HIV-1) transmission. Extending CAB dosing to a yearly injectable can advance efforts leading to the elimination of viral transmission. The current submission adds rigor, reproducibility and mechanistic insights for the extended apparent half-life of a yearlong antiretroviral injectable. Pharmacokinetic (PK) profiles of a nanoformulated fatty acid ester CAB prodrug (named NM2CAB) were affirmed at two academic and one contract research laboratory. PK profiles showed plasma CAB levels at or above the protein-adjusted 90% inhibitory concentration for up to one year after a single dose. Measures of drug biodistribution demonstrated sustained native drug at the muscle injection site and in lymphoid tissues (spleen and lymph nodes). The results paralleled NM2CAB uptake and retention in human macrophages. NM2CAB nanocrystals were stable in blood, tissue and liver homogenates. The long apparent drug half-life followed pH-dependent slow prodrug release in weeks from the nanocrystal. In contrast, solubilized prodrug was hydrolyzed in hours in plasma and tissues recorded from multiple mammalian species at basic pH. No measurable toxicities were recorded. These results, taken together, affirm the pharmacological mechanistic properties of a year-long nanoformulated CAB prodrug supporting the established protocol design for formulation safety, rigor and reproducibility.

---

While wide-spread availability of antiretroviral therapy (ART) has reduced human immunodeficiency virus type one (HIV-1) morbidities and mortality, viral transmission continues to persist. This is highlighted by yearly recordings of two million new infections worldwide ^1^. ART requires life-long daily oral regimens with both short and long-term toxicities. Changing dosing regimens driven by the emergence of resistant viral strains and regimen adherence are therapeutic limitations^2,3,4,5^. A priority for human immunodeficiency virus type one (HIV-1)/acquired immune deficiency syndrome (AIDS) research is how best to prevent the spread of infection which can be achieved through ART-mandated pre-exposure prophylaxis (PrEP)^6^. PrEP success has been buoyed by a long-acting (LA) parenteral formulation of cabotegravir. This permits every other month antiretroviral drug (ARV) dosing. CAB LA demonstrates improved pharmacokinetic and pharmacodynamic (PK and PD) profiles compared to more conventional ARV regimens and as such provides a promising new strategy for HIV-1 prevention and treatment^7^.

CAB is unique in its structural and antiretroviral properties. CAB LA is the first long acting injectable regimen amongst the group of integrase strand transfer inhibitors (INSTI). These relatively new ARVs have complemented existing ARVs based on their unique treatment and improved PK profiles. Notably, CAB LA PK results are highlighted by an extended plasma half-life of up to 54 days ^8^ and limited drug-drug interactions (DDI). The latter is linked both to its membrane permeability and low affinity for cytochrome P450 (CYP450)^9, 10^. Moreover, CAB has low aqueous solubility and a high melting point allowing its administration as a high concentration parenteral formulation^11^. CAB LA is currently completing phase 3 clinical trials and was approved recently for use in Canada. USA Food and Drug Administration (FDA) approval is anticipated in late spring-summer of 2021^12^. Nonetheless, there are limitations for CAB LA as the current formulation requires a 2 ml dosing volume known to elicit injection site reactions and continuous health care oversights^13^. Thus, newer formulations with longer dosing intervals, ease of access and reduced administration volumes will positively impact wide-spread regimen use.

In order to overcome the treatment limitations of CAB LA we developed lipophilic fatty acid ester CAB prodrugs nanoformulations (named NMCAB and NM2CAB, catalogued based on carbon lengths) with extended PK properties. The PK profiles were improved from months (NMCAB)^14^ to one year (NM2CAB)^15^. The early development of a year-long CAB, however, has met with a number of questions linked to reproducibility in different animal species and the underlying prodrug hydrolysis mechanisms to produce the observed extended drug half-life remained unclear. Moreover, safety measurements for the year-long parenteral formulation were required. With these in mind, we now report, from three separate laboratories, PK and biodistribution (BD) profiles, of NM2CAB, our lead nanoformulated stearoylated prodrug. Work was completed in rodents with head-to-head comparisons against a CAB LA formulation (reflected by NCAB) analogous to that currently being evaluated in human trials. Rigor and reproducibility are defined by PK studies by two research (from the University of Nebraska Medical Center Colleges of Medicine and Pharmacy) and one contract laboratory (Covance Laboratories, a global contract research organization and drug development services company) employing multiple mouse strains (immune deficient and competent) and rats. Lastly, mechanisms were elucidated together with the nanoformulation stability demonstrating that particle stability and slow prodrug dissolution best reflected the extended drug half-life for the prodrug formulation more than the enzymatic or chemical hydrolysis rates. Each of these tests proved critical in defining the safety and reproducibility of a year-long CAB apparent half-life. The solid state form of M2CAB nanocrystals is a key determinant for prodrug stability in biological fluids and tissues. We posit that translation of a safe biocompatible formulation will have a significant impact on the therapeutic armamentarium in preventing HIV/AIDS infection and in providing improved access to those populations most impacted by high viral transmission rates.

## Results

### Rigor and Reproducibility

The focus of the experiment methods and results were to ensure rigor into a new concept for ARV delivery. This concept centered on the change in parenteral drug administration from a once a month or every other month into a once a year drug administration therapeutic regimen. If substantiated, we posit that such an extension in drug usage would ease adherence to PrEP treatments and as a consequence serve to reduce transmission rates for HIV/AIDS and most notably in resource limited settings. Thus, the results outlined served to affirm prior published data sets by extending the analyses to additional mammalian species and serve to further substantiate each of the data sets by defining the mechanisms for extended half-life of the nanoformulated CAB prodrug.

### PK Determinations

In the first phase of these experiments, we evaluated the drug and prodrug PK and BD profiles in male Balb/c mice and Sprague Dawley (SD) rats that were injected intramuscularly (IM) with a single NCAB or NM2CAB dose of 45 mg CAB equivalents/kg. In mice, the peak CAB plasma concentration (C_max_ = 49,066 ng/ml) after NCAB administration was 6.5-fold higher than that of NM2CAB (C_max_ = 7,480 ng/ml) and occurred after 24 h (T_max_) for both formulations (**Table 1**). Following these recorded drug levels, CAB blood concentrations fell sharply for NCAB compared to NM2CAB. CAB terminal half-life (t_0.5_) following NM2CAB (118.4 days) treatment was 16-fold greater than NCAB (7.5 days). Similarly, CAB mean residence time (MRT) after NM2CAB administration was 19-fold longer than that of NCAB (164.7 vs. 8.7 days, respectively). The longer CAB t_0.5_ associated with NM2CAB is the result of a 20-fold higher apparent volume of distribution (Vz = 21.7 L/kg vs. 1.08 L/kg for NCAB). Starting at day 28 after NM2CAB administration, CAB plasma concentrations exceeded that following NCAB treatment at all time points, and by day 364 it was 280 times higher (263.2 ng/ml from NM2CAB vs. 0.9 ng/ml from NCAB). CAB plasma concentrations after NM2CAB administration remained higher than the CAB protein-adjusted 90% inhibitory concentration (PA-IC_90_) of 166 ng/ml^10^ for the entire period of the study, compared to less than 42 days following NCAB administration (**Figure 1**). M2CAB prodrug concentrations in plasma and blood (**Supplementary Figure 1**) after day 1 were more than 100 times lower than CAB concentrations or undetectable after NM2CAB administration. The peak plasma concentration of M2CAB (C_max_ = 558 ng/ml) was detected immediately after NM2CAB administration at 4 h (T_max_). After that, M2CAB concentrations declined rapidly and were undetected after day 42.

**Table 1.** CAB PK parameters after NCAB and NM2CAB administrations*.

| PK parameters | Mouse |  | Rat |  | Monkey |
| --- | --- | --- | --- | --- | --- |
|  | NCAB | NM2CAB | NCAB | NM2CAB | NM2CAB |
| $T_{max}$ (day) | 1.0 | 1.0 | 7.0 | 14.0 | 1.0 |
| $C_{max}$ (ng/ml) | 49,066.7 | 7,480.0 | 77,700.0 | 14,123.3 | 2,753.0 |
| $\lambda_z$ (1/h) | 0.0927 | 0.0059 | 0.0443 | 0.0085 | 0.0038 |
| $t_{0.5}$ (day) | 7.5 | 118.4 | 15.7 | 81.5 | 184.5 |
| $AUC_{last}$ (day*ng/ml) | 450,906.3 | 309,193.1 | 1,850,671.2 | 1,466,869.0 | 74,955.9 |
| $AUC_{0-\infty}$ (day*ng/ml) | 450,916.4 | 354,157.1 | 1,850,738.1 | 1,545,319.2 | 79,919.1 |
| $AUC_{\%}$ Extrapolation | 0.2% | 12.70 | 0.004 | 5.08 | 6.21 |
| $V_z/F$ (L/kg) | 1.08 | 21.70 | 0.55 | 3.42 | 149.84 |
| $CL/F$ (L/day/kg) | 0.100 | 0.127 | 0.024 | 0.029 | 0.563 |
| $MRT_{0-\infty}$ (day) | 8.8 | 164.8 | 21.6 | 117.7 | 228.4 |

\*PK parameters of CAB in Balb/c mice and Sprague-Dawley (SD) rats after NCAB and NM2CAB administration and in rhesus macaques after NM2CAB administration. PK parameters were calculated by non-compartmental analysis from the mean of N=6 mice and rats, and N=4 monkeys.

**Figure 1.**
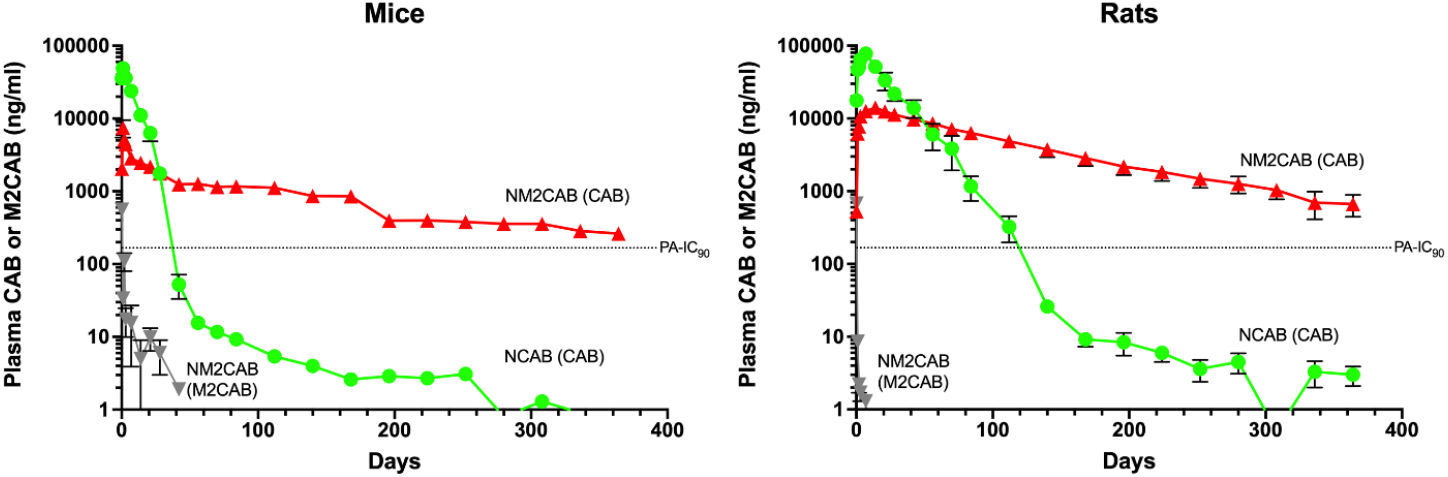
Year-long measurements of CAB and M2CAB plasma concentrations in mice and rats. Plasma concentration over time were recorded for CAB and M2CAB over one year after a single IM injection (45 mg CAB equivalent/kg) of NCAB or NM2CAB in male Balb/cJ mice (left-panel) and male SD rats (rightpanel). Data are expressed as mean ± SEM (N=6).

In rats, the peak plasma concentration (C_max_ = 77,700 ng/ml) after NCAB administration was 5.5 times higher than that of NM2CAB (C_max_ = 14,123 ng/ml) and occurred after 7 and 14 days (T_max_) for NCAB and NM2CAB, respectively (**Table 1**). Similar to mice, CAB plasma concentrations fell sharply and at a much faster rate after NCAB administration compared to NM2CAB. CAB t_0.5_ following NM2CAB (81.5 days) treatment was 5.2-fold greater than NCAB (15.7 days). Similarly, CAB MRT of NM2CAB was 5.5-fold longer than NCAB (117.7 vs. 21.6 days, respectively). The longer CAB t_0.5_ associated with NM2CAB is also the result of a 6.2-fold higher apparent volume of distribution (Vz = 3.42 L/kg vs. 0.55 L/kg for NCAB). Starting at day 56 after NM2CAB administration, CAB plasma concentrations exceeded that of NCAB at all-time points, and by the end of the study (day 364) were 225-fold higher (667.6 ng/ml from NM2CAB vs. 3.0 ng/ml from NCAB). CAB plasma concentrations after NM2CAB administration remained higher than the CAB PA-IC_90_ (166 ng/ml)^10^ for the entire period of the study, compared to less than 140 days after NCAB administration (**Figure 1**). The peak plasma concentration of M2CAB (C_max_ = 677 ng/ml) was detected immediately after NM2CAB administration at 4 h (T_max_). After that, M2CAB concentrations in plasma and blood (**Supplementary Figure 1**) sharply decreased and were undetected after day 7. In follow-on we extended the PK analyses in rhesus macaques (RM) from what was previously published for NM2CAB ^15^. In these extended studies the PK profiles of NM2CAB were evaluated for up to 2 years. Animals were injected IM with a single dose of 45 mg CAB equivalents/kg of NM2CAB. Similar to mice and rats, NM2CAB was slowly eliminated from blood and CAB was detected for the entire period of the study (data not shown). After that, CAB plasma concentrations decreased slowly over the period of ~ 2 years (t_0.5_ = 184 days and MRT = 228 days) (**Table1**).

We sought to confirm these PK test results in studies conducted by an independent facility. For these experiments blinded samples of NCAB and NM2CAB formulations were sent to Covance, a contract research laboratory. These experiments were designed to provide rigor and reproducibility of the enhanced PK profiles for NM2CAB. PK tests were performed in two mouse strains, male and female, for NCAB and NM2CAB at two CAB equivalent doses for each formulation. As shown in **Figure 2A-D**, results from both mouse studies at doses of 45 or 70 mg CAB equivalents/kg showed that CAB plasma concentrations after NCAB administration were initially higher compared to NM2CAB. After 28 days, CAB concentrations fell sharply and at a much faster rate after NCAB compared to NM2CAB treatment after either dose. In both strains of mice given the lower dose of NCAB, CAB was below 10 ng/ml at three months, and at the limit of detection (1 ng/ml) after four months (**Figure 2A, B**). In contrast, after NM2CAB administration CAB concentration decreased at a much slower rate and at 6 months was 437 and 575 ng/ml in male Balb/c and female NSG mice, respectively. The higher dose (70 mg CAB equivalents/kg) provided a 2.2- to 2.8-fold increase in plasma CAB AUC compared to the lower dose in male Balb/c and female NSG mice, respectively (**Figure 2C, D**). In addition, samples from the study were analyzed at UNMC using a different bioanalytical method and provided values that were similar to those obtained by the Covance analyses (**Figure 2E, F; Supplementary Figure 2**).

**Figure 2.**
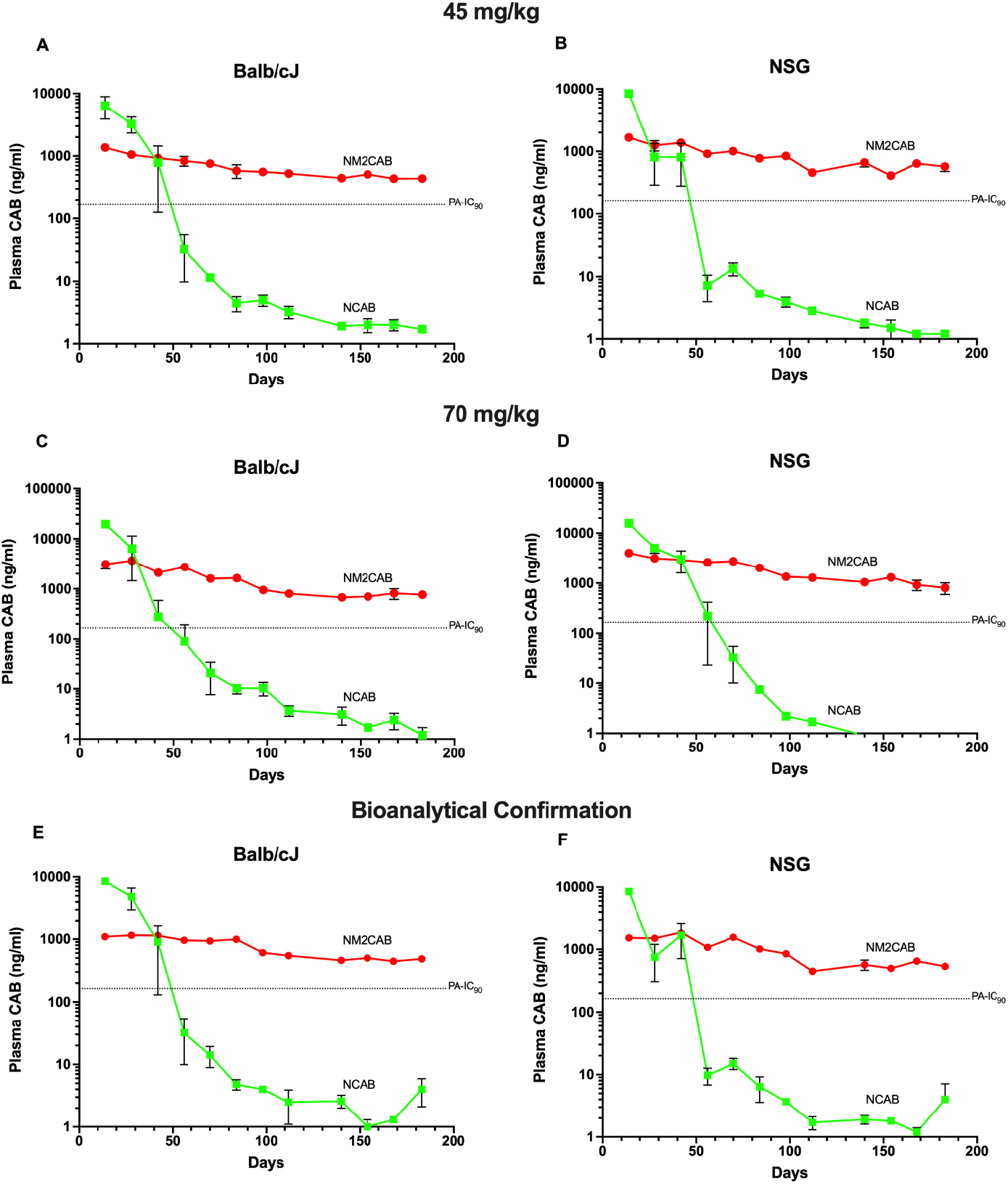
CAB plasma concentrations in Balb/cJ and NSG mice. Male Balb/cJ or female NSG mice were given a single intramuscular injection of 45 or 70 mg CAB equivalents/kg of NCAB or NM2CAB. Plasma was collected every other week for 6 months at which time the study was terminated. CAB concentrations were quantitated by LC-MS/MS by Covance laboratories (A-D); treatment with 45 mg/kg in (A) male Balb/cJ and (B) female NSG mice; treatment with 70 mg/kg in (C) male Balb/cJ and (D) female NSG mice. CAB concentrations in plasma samples were also quantitated at UNMC (bioanalytical confirmation). Plasma CAB concentrations for the 45 mg/kg treatment groups are shown for (E) male Balb/cJ and (F) female NSG mice. Data are expressed as mean ± SEM (N=2-6 animals per group per timepoint).

## Mechanisms for the Extended CAB Half-Life

### Prodrug Stability and Drug Release

To assess the stability of the prepared prodrug nanoformulation, the prodrug and parent drug contents of the suspension were determined over time at different suspension dilutions and in various biological matrices. Both neat and 20X-diluted nanoformulations demonstrated storage stability and equivalent release kinetics when assessed at room temperature. The neat nanoformulations are prepared for each of the animal studies while dilutions are made for cell-based testing of uptake, retention, and release. NM2CAB demonstrated sustained nanoparticle and prodrug stability with minimal drug release from the nanoparticles during storage for one year compared to NCAB. We detected an up to 8% release of M2CAB from NM2CAB at day 0, without subsequent drug release for one year. During the observation period, NM2CAB was completely stable (**Table 2**). Nanoformulations of >300 mg/ml prodrug content were manufactured with no loss of particle integrity. In contrast, under the same storage conditions, a burst release of 35% was recorded for NCAB.

**Table 2.** Storage stability of NM2CAB and NCAB nanoformulations*.

|  | Time (Day) | NCAB |  | NM2ACB |  |
| --- | --- | --- | --- | --- | --- |
|  |  | Stability | % Release | Stability | % Release |
| Undiluted | 0 | 90.0% | 35.8% | 112.3% | 7.4% |
|  | 4 | 88.9% | 38.4% | 95.2% | 6.7% |
|  | 7 | 103.2% | 30.0% | 83.6% | 10.8% |
|  | 14 | 102.4% | 26.0% | 92.7% | 7.0% |
|  | 30 | 100.5% | 24.1% | 95.8% | 8.1% |
|  | 60 | 95.8% | 39.3% | 108.4% | 6.2% |
|  | 90 | 95.3% | 32.5% | 94.3% | 8.4% |
|  | 252 | 95.9% | 37.6% | 112.2% | 6.3% |
|  | 365 | 111.7% | 26.4% | 100.8% | 5.0% |
| 20X-diluted | 0 | 110.4% | 29.8% | 94.8% | 7.9% |
|  | 4 | 106.5% | 26.5% | 84.3% | 9.7% |
|  | 7 | 101.5% | 30.8% | 87.1% | 8.6% |
|  | 14 | 111.3% | 23.7% | 106.8% | 6.1% |
|  | 30 | 101.4% | 25.0% | 86.9% | 8.6% |
|  | 60 | 114.8% | 20.3% | 116.1% | 7.0% |
|  | 90 | 105.5% | 22.1% | 105.6% | 7.4% |
|  | 252 | 102.7% | 31.1% | 102.3% | 8.5% |
|  | 365 | 101.6% | 36.2% | 102.3% | 5.4% |

\*Stability of NM2CAB and NCAB at 55.4 and 36.7 mg/ml (neat) and 2.77 and 1.835 mg/ml (20X diluted), respectively.

Equivalent stability and release kinetics were seen in both 20X-diluted and undiluted (neat) formulations (**Table 2**). Next, we assessed chemical stability of the solubilized form of M2CAB at a range of pHs [pH 1.0 (0.1M HCl), pH 7.4 (PBS), pH 8.3 (heat inactivated plasma) and pH 11.0 (0.1 M NaOH)]. All were tested at 37°C over 24 h. At 24 h, percent M2CAB recovered was 87, 52, 5, and 2 at pH 1.0, pH 7.4, pH 8.3 and pH 11.0, respectively (**Supplementary Figure 3**). Reductions in M2CAB prodrug paralleled native CAB formation. The results highlighted and affirmed the direct links between nanocrystal dissociation, prodrug solubility and chemical stability. The data underlies the potential importance of controlling prodrug solubility within the formulation to affect hydrolysis rates and apparent drug half-lives.

### Solubilized Stable Prodrug and Nanoformulation

In order to evaluate chemical stabilities of the compounds in solid state and solubilized forms, a solution of M2CAB and NM2CAB and NCAB nanocrystals were evaluated in mouse, rat, rabbit, monkey, dog, and human blood over 24 h (**Figure 3; Supplementary Figure 4**). The solubilized prodrug (M2CAB solution) was unstable in all species. Greater than 80% of M2CAB was hydrolyzed after 6 h in dog, monkey and human (**Figure 3A, C**). M2CAB was unstable in heat-inactivated plasma (**Figure 3E**), indicating that its degradation in blood was due to chemical instability independent of a biological matrix. The decline in M2CAB prodrug paralleled formation of native CAB (**Figure 3B, 3D, 3F**). In contrast, NM2CAB was stable in blood from all species with > 80% of M2CAB recovered and < 10% of CAB formed after 24 h (**Figure 3A, C**), suggesting that nanoparticle stability and M2CAB solid forms within NM2CAB nanocrystals controls prodrug dissolution. Similarly, both CAB and NCAB were stable for 24 h in blood in all species (**Supplementary Figure 4**). The metabolic stability of M2CAB solution was next assessed in liver S9 fractions from mouse, rat, rabbit, monkey, dog and humans (**Supplementary Figure 5**). In contrast to blood, < 25% of M2CAB was lost in a 2 h incubation and was accounted for by CAB formation. No species-specific differences were observed. Similar to blood, chemical, rather than biological factors, accounted for the instability. These reactions were next assessed in rat liver, spleen, muscle, lymph node, and heat-inactivated liver homogenate (**Supplementary Figure 6**). In rat tissues, 50-80% of M2CAB solution remained after 6 h of incubation. M2CAB solution was also unstable in the control heat-inactivated liver homogenate, indicating that its degradation is due to chemical instability upon solubilization, and is independent of the biological matrices under similar assay conditions. The decline in M2CAB prodrug was accounted for by formation of CAB in all tissues (**Supplementary Figure 6**). In contrast to solubilized M2CAB, the prodrug within NM2CAB solid drug nanoparticles was stable in all tissues with 88% of M2CAB recovered and 10% of CAB formed after a 6 h incubation (**Supplementary Figure 6**).

**Figure 3.**
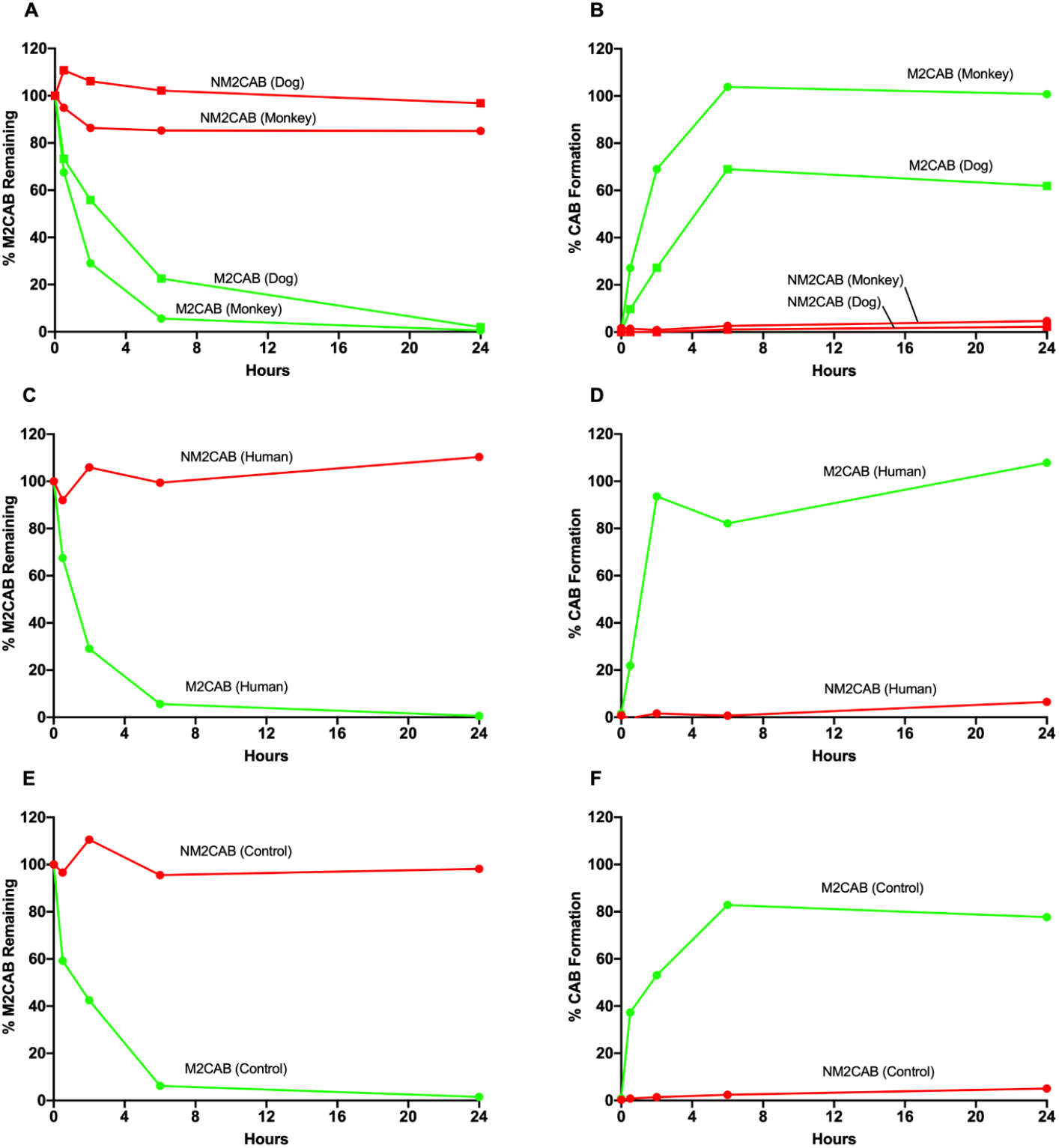
Metabolic CAB release-stability profiles during prodrug hydrolysis. Stability profiles of prodrug (M2CAB) and nanoformulated prodrug (NM2CAB), native CAB, and nanoformulated CAB (NCAB, reflective of current phase III trials) were used at 1 µM concentration and drug concentrations determined in blood from three mammalian species (monkey, dog, and human) and in heat-inactivated plasma as a control. Data shown as M2CAB (disappearance for M2CAB over time (alone and from NM2CAB nanoformulations) in (A) monkey and dog (B) human, and (E) heat-inactivated plasma. CAB appearance over time recorded from M2CAB and NM2CAB in (B) monkey and dog, (D) human, and (F) heat-inactivated plasma (control) appear in parallel measurements.

### Cellular Uptake, Retention and Prodrug Release

With the stability of the prodrug nanoformulation affirmed we next evaluated the efficiency of the prodrug nanoformulation for uptake, retention and release rates in macrophages. Macrophages represent the primary cell type known to store the particles and the efficiency of uptake reflects the established depot of drug during the extended times observed *in vivo^15^*. To these ends, human monocyte-derived macrophages (MDMs) were used to record cell uptake, retention, and release of native-CAB, M2CAB, NCAB, and NM2CAB (**Figure 4**). Up to 60, 30, 6 and 0.1% of NM2CAB, M2CAB, NCAB, CAB were taken by MDMs at 8 h, respectively (**Figure 4A**). After cells were loaded for 8 h with each of the formulations they were evaluated for up to 30 days to determine drug retention and release. Seventy-five percent and 4% of intracellular NM2CAB and M2CAB were retained by MDMs at one month, respectively. Notably, intracellular CAB levels for NCAB and native CAB were below or at the limit of detection within hours of drug loading. M2CAB and CAB release from cells was determined by their appearance in culture media (**Figure 4B**). Ninety-nine percent of the retained intracellular NM2CAB was in the form of the M2CAB prodrug, while 80% of the released extracellular NM2CAB was in the form of the parent CAB. We also tested stability and CAB formation for both NM2CAB and M2CAB in the macrophage cell culture media and in MDM-cell lysate (as a surrogate for intracellular stability) over 15 days. NM2CAB nanoparticles were greater than 80% stable, while M2CAB solution was completely hydrolyzed into CAB both intracellularly and in the media (data not shown). The data sets demonstrate the importance of assaying primary macrophages as reservoirs for prodrug nanoformulations highlighting the retention and release of both drug (CAB) and prodrug (M2CAB) in these cell sites.

**Figure 4.**
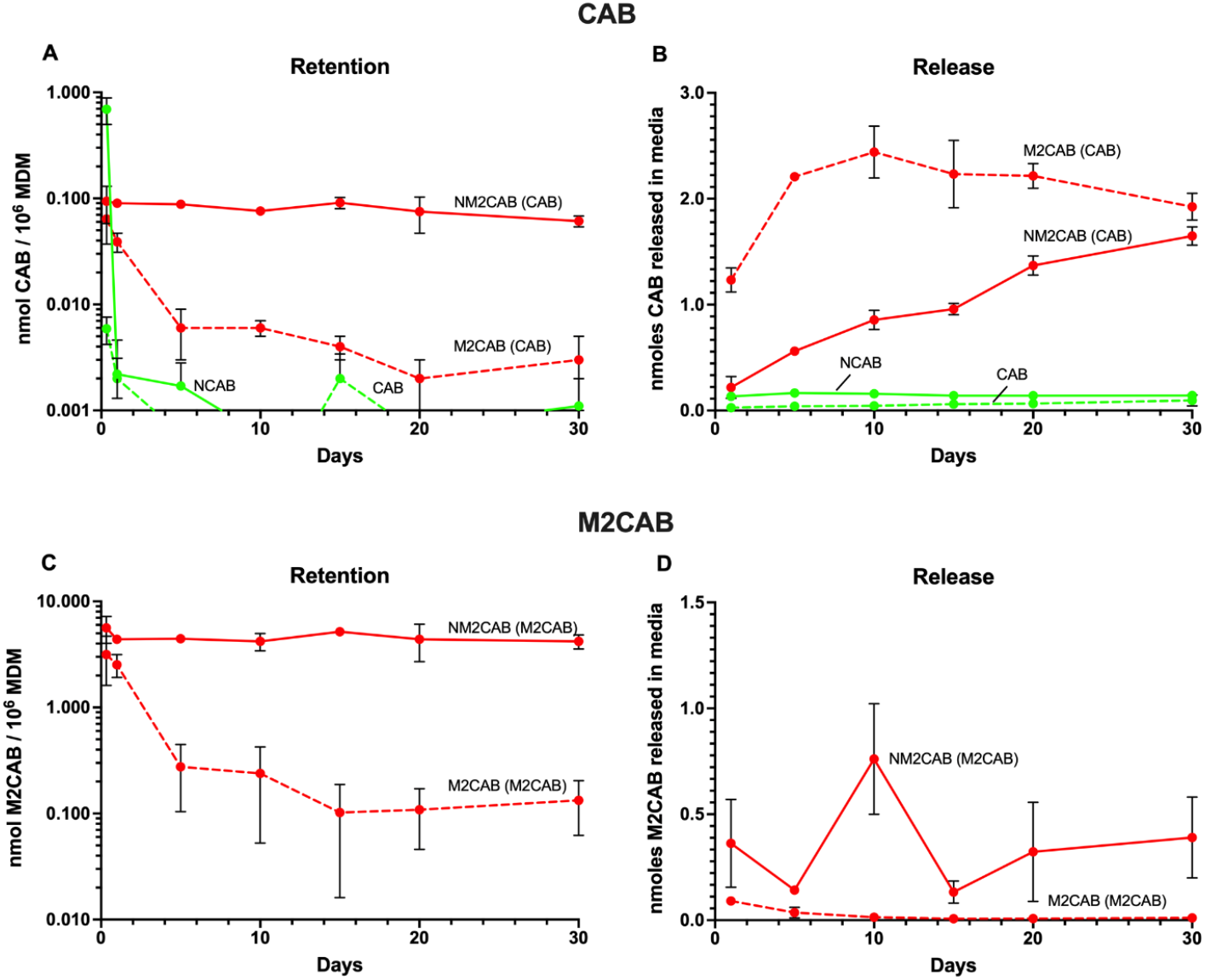
Cellular uptake, retention and release of native drug and prodrug. (A) Intracellular CAB, (B) CAB released in media, (C) Intracellular M2CAB, and (D) M2CAB released in media after native-CAB, native-M2CAB, NCAB, and NM2CAB treatment to MDMs at 10 µM (10 nmoles/10^6^ cells) for 8 h. Data are expressed as mean ± SD (N=3).

### Blood Drug Measures

Due to the instability of the solubilized form of M2CAB in blood, conditions were optimized to prevent *ex vivo* prodrug hydrolysis and CAB formation after sample collection. Methanol (containing 0.1% FA and 2.5 nM AF) inhibited the hydrolysis of prodrug in blood. Therefore, we collected blood directly into tubes containing 0.1% FA and 2.5 nM AF in methanol to prevent any *ex vivo* M2CAB hydrolysis into CAB. A portion of the collected blood samples was saved to produce plasma. CAB blood concentrations were nearly 2-fold lower than that in plasma in all samples, and this was also shown previously ^16, 17^ (**Supplementary Figure 1**). Because blood concentrations of the prodrug were extremely low (< 1% of CAB), plasma samples seemed to be stabilized by EDTA, and because of the low-temperature storage (−20°C), M2CAB blood and plasma concentrations were very similar. Therefore, both plasma and blood sample collections are valid for M2CAB analysis and there was no *ex vivo* hydrolysis of prodrug or parent drug formation during sample preparation, storage, and analysis.

### Tissue CAB Distribution

After both NM2CAB and NCAB treatment, CAB was widely distributed to all tissues with the exception of brain in both mice and rats. The highest CAB and M2CAB concentrations were at the site of injection and in lymph nodes, followed by spleen, lung, kidney, and liver (**Table 3**). M2CAB concentrations in many tissues were higher than that of CAB at all time points. At day 28, CAB tissue concentrations in mice were ~16,000, 3,000, and 2,200 ng/g in the site of injection, lymph nodes, and spleen, respectively, while CAB concentrations were in the range of 100-250 ng/g in other tissues (liver, kidney, lung, muscle, gut) and 16 ng/g in the brain. Similarly, M2CAB concentrations at day 28 were 1.7×10^6^, 230,000, 35,000, and 3000 ng/g at the site of injection, lymph nodes, spleen, and liver, respectively, while in all other tissues it was in the range of 10-100 ng/g. By day 364 of NM2CAB administration in mice, CAB concentrations were 4,939 and 86 ng/g at the site of injection and lymph nodes, respectively, while M2CAB concentrations in the same tissues were 419,000 and 4,187 ng/g. The same trend in tissue distribution was observed after NCAB administration in mice, but with much lower CAB concentrations. By day 364, CAB concentration for NCAB at the site of injection was ~30 ng/g and undetected in all other tissues. Similarly, in rats the NM2CAB highest distribution was at the site of injection and lymph nodes. At day 28, CAB tissue concentrations were ~7,000, 2,200, and 1,800 ng/g in the site of injection, lymph nodes, and lung, respectively, while CAB concentrations were in the range of 400-800 ng/g in other tissues (liver, spleen, kidney, muscle, gut) and 97 ng/g in the brain. Similarly, M2CAB concentrations at day 28 were 7.6 ×10^6^, 458, and 339 ng/g at the site of injection, liver, and spleen, respectively, while in all other tissues were < 40 ng/g. By day 364 after NM2CAB administration in rats, CAB concentrations were 1,800 and 118 ng/g in the site of injection and lymph nodes, respectively, while M2CAB concentrations in the same tissues were 103,000 and 1.2 ng/g. Brain showed the lowest tissue distribution in both species after both NM2CAB and NCAB administration. By day 168, CAB brain concentrations were barely detectable after NCAB administration in both mice and rats, while NM2CAB administration produced ~8 and 44 ng/g in mice and rats, respectively. Also, the prodrug (M2CAB) brain concentrations were below detection limit in both mice and rats at all time points. CAB tissue concentrations were not only higher after NM2CAB compared to NCAB, but they also decreased at a much slower rate over time. Day 168 concentrations were 50 and < 1% on day 28 after NM2CAB and NCAB administration, respectively (**Table 3**). The summary PK and BD results are provided in graphic form that serves as a comparison between NCAB and NM2CAB (**Figure 5**). The figure provides, in cartoon form, the actual stability of the CAB prodrug nanoformulations when compared to native drug. This is done by illustrating sustained drug depots (pictured in green) at the injection site (deltoid muscle) and within the lymphoid system (lymph node and spleen) and liver for time periods measured at up to one year.

**Table 3.** CAB and M2CAB tissue distribution*.

| Animal | Time (Day) | Liver |  | Spleen |  | Kidney |  | Brain |  | Lung |  | Muscle |  | Gut |  | Lymph Node |  | Site of Injection |  |
| --- | --- | --- | --- | --- | --- | --- | --- | --- | --- | --- | --- | --- | --- | --- | --- | --- | --- | --- | --- |
|  |  | Mean (ng/g) | SEM | Mean (ng/g) | SEM | Mean (ng/g) | SEM | Mean (ng/g) | SEM | Mean (ng/g) | SEM | Mean (ng/g) | SEM | Mean (ng/g) | SEM | Mean (ng/g) | SEM | Mean (ng/g) | SEM |
| Mouse | **CAB concentrations from NCAB** |  |  |  |  |  |  |  |  |  |  |  |  |  |  |  |  |  |  |
|  | Day 28 | 76.7 | 10.6 | 253.3 | 58.1 | 217.2 | 49.6 | 14.6 | 3.9 | 753.8 | 257.5 | 123.3 | 30.0 | 149.4 | 34.7 | 621 | 150 | 16,437 | 7,127 |
|  | Day 84 | 0.6 | 0.3 | 1.0 | 0.2 | 2.1 | 0.7 | 0.0 | 0.0 | 3.0 | 0.2 | 0.5 | 0.1 | 1.7 | 0.9 | 15 | 5 | 306 | 50 |
|  | Day 168 | 0.0 | 0.0 | 0.1 | 0.1 | 0.4 | 0.1 | 0.0 | 0.0 | 4.8 | 0.9 | 0.6 | 0.3 | 0.5 | 0.3 | 5 | 2 | 169 | 11 |
|  | Day 364 | 0.0 | 0.0 | 0.0 | 0.0 | 0.4 | 0.3 | 0.0 | 0.0 | 0.0 | 0.0 | 0.4 | 0.3 | 0.0 | 0.0 | 0 | 0 | 30 | 4 |
|  | **CAB concentrations from NM2CAB** |  |  |  |  |  |  |  |  |  |  |  |  |  |  |  |  |  |  |
|  | Day 28 | 159.3 | 30.7 | 2,201.3 | 202.4 | 196.6 | 15.0 | 16.4 | 2.4 | 224.9 | 21.4 | 103.9 | 12.7 | 158.2 | 19.4 | 2,995 | 964 | 16,558 | 2,490 |
|  | Day 84 | 76.3 | 12.5 | 1,268.5 | 466.8 | 133.3 | 17.4 | 10.5 | 1.6 | 254.9 | 82.2 | 56.8 | 7.5 | 88.0 | 6.2 | 3,394 | 871 | 9,946 | 1,675 |
|  | Day 168 | 44.9 | 2.0 | 59.7 | 6.8 | 148.4 | 61.5 | 8.1 | 1.3 | 132.5 | 14.9 | 36.0 | 4.8 | 56.7 | 3.6 | 1,580 | 568 | 9,749 | 621 |
|  | Day 364 | 14.9 | 0.5 | 16.4 | 1.2 | 36.1 | 3.5 | 0.5 | 0.2 | 48.0 | 3.8 | 11.0 | 1.7 | 20.4 | 2.7 | 86 | 20 | 4,939 | 708 |
|  | **MCAB concentrations from NM2CAB** |  |  |  |  |  |  |  |  |  |  |  |  |  |  |  |  |  |  |
|  | Day 28 | 3,208.7 | 827.1 | 35,933 | 8,914 | 11.4 | 3.4 | 4.5 | 1.8 | 137.1 | 38.5 | 8.9 | 4.4 | 9.4 | 2.6 | 231,685 | 135,071 | 1,757,500 | 259,624 |
|  | Day 84 | 558.6 | 271.4 | 13,438 | 4,543 | 1.7 | 0.5 | 0.5 | 0.1 | 53.7 | 26.4 | 8.8 | 3.8 | 3.7 | 1.5 | 274,090 | 98,711 | 741,500 | 175,267 |
|  | Day 168 | 39.8 | 6.9 | 492 | 50 | 1.1 | 0.3 | 5.8 | 1.9 | 12.7 | 3.8 | 1.2 | 0.4 | 2.4 | 0.8 | 81,752 | 33,916 | 756,333 | 83,539 |
|  | Day 364 | 5.3 | 0.7 | 33 | 4 | 2.2 | 1.4 | 0.0 | 0.0 | 3.5 | 2.3 | 1.8 | 1.4 | 3.3 | 1.9 | 4,188 | 1,550 | 419,000 | 36,491 |
| Rat | **CAB concentrations from NCAB** |  |  |  |  |  |  |  |  |  |  |  |  |  |  |  |  |  |  |
|  | Day 28 | 1,411.2 | 296.5 | 855.1 | 206.0 | 2,351.6 | 624.3 | 216.0 | 54.1 | 2,747.8 | 667.9 | 3,071.2 | 443.3 | 1,086.0 | 262.5 | 3,105 | 488 | 675,486 | 250,918 |
|  | Day 84 | 55.6 | 15.4 | 124.2 | 39.4 | 164.4 | 35.1 | 13.3 | 4.5 | 420.4 | 203.0 | 300.9 | 74.6 | 127.4 | 44.1 | 485 | 182 | 1,219 | 956 |
|  | Day 168 | 0.6 | 0.2 | 3.0 | 1.0 | 39.8 | 5.1 | 1.6 | 0.9 | 3.7 | 0.8 | 6.6 | 1.9 | 9.6 | 3.3 | 7 | 1 | 43 | 28 |
|  | Day 364 | 0.0 | 0.0 | 0.0 | 0.0 | 0.5 | 0.2 | 0.0 | 0.0 | 2.5 | 1.1 | 0.0 | 0.0 | 0.1 | 0.0 | 0 | 0 | 16 | 1 |
|  | **CAB concentrations from NM2CAB** |  |  |  |  |  |  |  |  |  |  |  |  |  |  |  |  |  |  |
|  | Day 28 | 674.1 | 184.6 | 434.1 | 84.3 | 779.0 | 158.1 | 97.7 | 19.2 | 1,824.5 | 375.4 | 560.5 | 110.5 | 692.7 | 126.8 | 2,210 | 401 | 6,821 | 693 |
|  | Day 84 | 331.0 | 61.2 | 473.2 | 68.9 | 552.7 | 78.7 | 74.3 | 15.3 | 1,490.7 | 253.5 | 506.6 | 112.1 | 636.8 | 128.9 | 1,411 | 224 | 4,893 | 990 |
|  | Day 168 | 177.0 | 53.8 | 126.3 | 27.7 | 323.9 | 70.5 | 43.7 | 10.0 | 334.1 | 52.1 | 228.6 | 67.2 | 175.7 | 31.2 | 723 | 171 | 1,915 | 449 |
|  | Day 364 | 44.4 | 14.0 | 44.2 | 16.4 | 152.0 | 41.4 | 5.4 | 2.6 | 149.4 | 61.8 | 26.4 | 9.9 | 54.5 | 18.1 | 119 | 49 | 1,795 | 261 |
|  | **MCAB concentrations from NM2CAB** |  |  |  |  |  |  |  |  |  |  |  |  |  |  |  |  |  |  |
|  | Day 28 | 458.1 | 66.9 | 339.3 | 48.3 | 1.9 | 0.2 | 0.5 | 0.1 | 37.5 | 28.3 | 27.8 | 18.1 | 35.8 | 10.5 | 12 | 3 | 762,500 | 72,235 |
|  | Day 84 | 264.4 | 90.1 | 302.0 | 58.4 | 0.7 | 0.1 | 0.3 | 0.1 | 24.6 | 4.9 | 0.8 | 0.2 | 4.3 | 0.6 | 9 | 1 | 240,566 | 89,748 |
|  | Day 168 | 51.9 | 9.9 | 89.2 | 22.5 | 1.3 | 0.2 | 0.6 | 0.1 | 5.9 | 1.2 | 0.6 | 0.2 | 2.5 | 0.5 | 8 | 2 | 108,000 | 40,875 |
|  | Day 364 | 7.9 | 2.4 | 8.8 | 2.8 | 2.7 | 2.7 | 0.0 | 0.0 | 0.4 | 0.4 | 0.6 | 0.3 | 0.0 | 0.0 | 1 | 0 | 102,937 | 15,663 |

\*CAB (native) and M2CAB (prodrug) tissue distribution in Balb/c mice and SD rats after a single IM injection of 45 mg CAB equivalents/kg NCAB and NM2CAB (N=6; Mean ± SEM).

**Figure 5.**
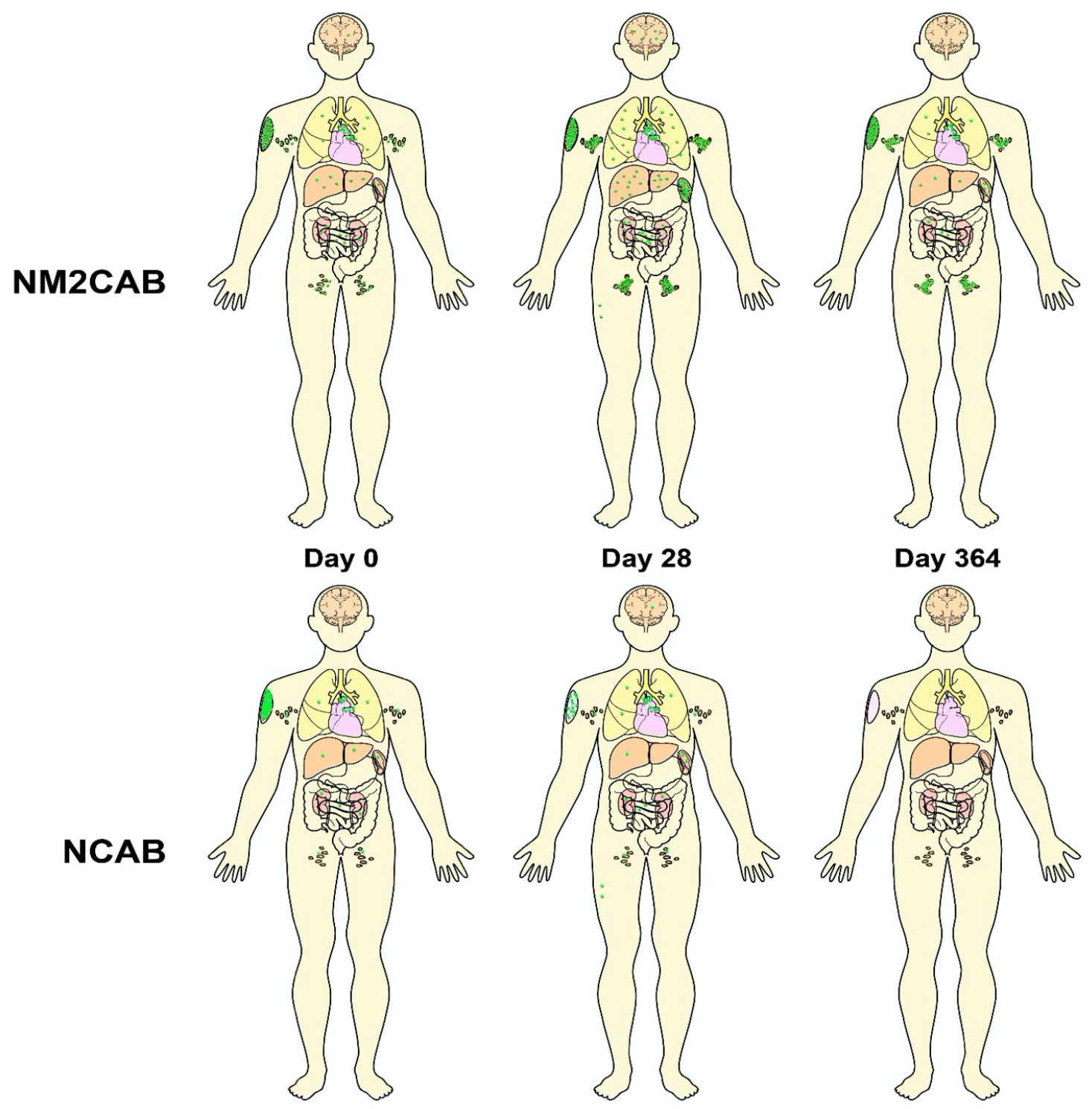
Pharmacokinetic and biodistribution (PK and BD) models of NM2CAB and NCAB formulation are illustrated. In the cartooned a human is injected intramuscularly with NM2CAB (top panels, a nanoformulated stearoylated fatty ester CAB prodrug) or NCAB (bottom panel, a nanoformulated native drug). PK and BD profiles showed that the nanocrystal prodrug of NM2CAB remained at high levels at the site of injection and throughout lymphoid organs (lymph nodes and spleen) for up to one year. Free native drug was released slowly and continuously in blood during that time. Particles dissolution occurs slowly as the drug is continuously present in circulation. This underlies the maintenance of CAB in plasma, at or above the protein-adjusted 90% inhibitory concentration for one year. The illustration serves to affirm the study’s conclusions that NM2CAB solid drug nanocrystals are stable in blood and tissue. NCAB, in contrast, is present at the site of the deltoid muscle injection and cleared more rapidly. The high-level distribution to the lymph nodes and spleen observed at one month are not detectable beyond that time period. Detectable CAB prodrug nanoformulations are observed in lymph nodes and spleen followed by liver, lungs, intestines and kidneys through a year of observation in NM2CAB injected hosts.

## Discussion

CAB is a second generation INSTI currently nearing USA FDA approval as a LA injectable ^18^. While effective in PrEP and has generated attention by the scientific and patient community for both treatment and prevention of HIV/AIDS ^12, 17, 19^ limitations in dosing volumes, intervals of administration and local injection site reactions remain. To overcome these, prodrug nanoformulated were developed and are currently undergoing pre-clinical evaluations by our group ^14, 15^. Early proof of concept studies showed encouraging results. However, what was done previously in support of NM2CAB required affirmation and safety evaluation to better position the product for future clinical testing. Underlying mechanisms linked to a year-long medicine also required evaluation. Towards both ends, independent laboratories employed two strains of mice and rats to affirm our prior works. Results showed sustained CAB plasma levels for at least one year in all species. Plasma concentrations were up to 100-fold higher than those resulting from NCAB (an equivalent preparation to the CAB LA). Most notably we now show that the mechanism of this extended half-life was attributed to prodrug solubility and release from the nanoformuation rather than drug hydrolysis (chemical and enzymatic) or tissue drug-particle biodistribution. These findings offer novel insights into how best to bring such extended release medicines to the clinic with broad relevance to ARVs now being modified as LA formulations.

INSTIs are a newer class of antiretrovirals (ARVs) which prevent the integration of viral DNA into the host cell DNA, by inhibiting the viral integrase enzyme ^20^. The main advantage of INSTIs rests in their activity against HIV mutant viruses resistant to other ARV classes ^21^. Raltegravir (RAL) was the first INSTI approved by the USA FDA in 2007; elvitegravir (EVG), dolutegravir (DTG), and bictegravir (BIC), were next. CAB is the newest addition to this class, and is currently under development by ViiV Healthcare, Research Triangle Park, NC, USA ^18^.

CAB has more favorable PK properties then other INSTIs. This includes its long plasma half-life of 30-40 h ^8^, fast oral absorption (t_max_ = 1.5-2 h) ^16^, near-complete oral bioavailability ^16, 22^, and low DDI potential due to its high membrane permeability and low affinity for CYP450 ^9, 10^. In humans, CAB is primarily eliminated unchanged in feces (up to 50% unchanged and 10% unidentified metabolites) and as glucuronide metabolites in urine (27%) ^16^. In addition, CAB is characterized by low aqueous solubility (0.015 mg/mL in water), and high melting point (248-251 °C), which permits its loading at high concentrations as nanosuspensions ^11^. CAB LA is in late stage clinical trials ^12^. When administered as a LA parenteral, CAB concentrations remain above four-times the PA-IC_90_ of 0.664 ug/ml, for 16 weeks after an 800 mg loading dose ^13^. With a drug load of 400 mg CAB can maintain plasma drug levels beyond four-times the PA-IC_90_ at 8 weeks ^23^. The combination of CAB and rilpivirine LA in infected adults (the LATTE-2, Long-Acting antiretroviral Treatment Enabling trial) showed that 400, 600 and 800 mg administered every 2, 8 and 12 weeks was as effective as once daily 30 mg CAB given orally. Therefore, newer formulations that allow longer dosing intervals and/or lower dosing volumes can markedly improve what is now available for CAB LA.

To this end, our group has recently developed several lipophilic fatty acid ester-CAB prodrugs (MCAB) prepared as long-acting nanoformulations. NMCAB and NM2CAB markedly improved and extended the PK profile of CAB compared to CAB LA ^15^. In this paper, we expanded on our earlier work by reporting the PK and BD of the lead nanoformulated CAB prodrug (NM2CAB) in mice and rats and compared it to NCAB, a formulation reflective of CAB LA. Our year-long NM2CAB was reproducible and robust as demonstrated by results from three independent laboratories in two strains of mice and in three animal species including mice, rats, and rhesus macaques. Also, we elucidated the absorption, distribution, metabolism, and excretion and mechanisms behind the enhanced PK profiles of NM2CAB using disposition measures of CAB’s pharmaceutical compound profiles within the tested organism and drug release kinetics.

In the current study, we characterized the metabolism, chemical and biological stability, release kinetics, as well as the PK profile of NM2CAB in comparison to NCAB. We used several means to investigate the mechanisms behind the enhanced PK profile of NM2CAB. First, the chemical stability and release kinetics of NM2CAB were assessed over one year. The original and diluted NM2CAB were stable with a minimal burst release of 8% at day 0 without any further release to one year. NCAB showed similar results, but with a higher burst release of 35%. Stability of M2CAB in solubilized form was pH-dependent, where degradation to CAB accelerated with increasing pH and was minimal at pH 1.0. The NM2CAB formulations produced were stable for 360 days with respect to size (nm), polydispersity (PDI) and surface charge (zeta potential) (**Supplementary Figure 7**)

M2CAB is an ester-linked prodrug, which is hydrolyzed in tissues and blood ^15, 24^, presumably by esterases and pH-dependent chemical hydrolysis upon release and solubilization of the lipophilic prodrug. Species differences in the distribution of these esterases have been previously reported ^24^. Therefore, we quantified the metabolic stability of CAB, M2CAB, NCAB, and NM2CAB in blood and liver S9 fractions from five animal species and humans. In blood, the ester bond of the solubilized form of M2CAB prodrug was rapidly hydrolyzed into the parent CAB within 6 h in all species. In addition, M2CAB solution was unstable in the control heat-inactivated plasma and in PBS, indicating chemical instability, as well. In contrast, NM2CAB solid drug nanocrystals, CAB, and NCAB were stable in blood and more than 90% of M2CAB (NM2CAB) or CAB (CAB, NCAB) was recovered by 24 h in all six species. M2CAB solution was also hydrolyzed in liver S9 fraction from all species, but at a lower rate than in the blood. In addition, the metabolic stability of NM2CAB was quantified in several tissue (liver, spleen, muscle, lymph node) homogenates from rats. In general, the hydrolysis rate of M2CAB was slower in tissues than in blood.

Our mechanistic studies were also performed in MDMs. Macrophages serve as a major depot for CAB prodrug nanoformulations while at the same time a principal HIV-1 reservoir ^25, 26, 27^. Therefore, we quantified the uptake, retention, and release kinetics of NM2CAB in MDMs. NM2CAB was efficiently taken up and retained for at least 30 days by MDMs. NM2CAB demonstrated up to 100-fold greater uptake and retention by MDMs compared to NCAB and unformulated CAB. While unformulated solution of M2CAB prodrug demonstrated enhanced cellular uptake, it was not retained inside the cells and greater than 95% was released into the media within days and was recovered as CAB. M2CAB solution hydrolyzed into CAB both intracellularly and in the media while NM2CAB nanoparticles were stable both intracellularly and in the media, and slowly released M2CAB over 30 days. Collectively, this data shows that NM2CAB solid drug nanocrystals enhance cellular drug uptake and extends intracellular drug retention due to the slow release and solubilization of M2CAB.

PK studies were performed in mice, rats, and rhesus macaques. A single dose of NM2CAB demonstrated substantial improvements in CAB PK profile, reflected by elevated and sustained plasma and tissue CAB concentrations compared to NCAB. CAB terminal t_0.5_ was extended from 7.5 to 118.4 days in mice and from 15.7 to 81.5 days in rats after NM2CAB administration compared to NCAB. CAB plasma levels were maintained above the PA-IC_90_ of 166 ng/ml for at least one year after a single dose of NM2CAB to both mice and rats. By the end of study, CAB plasma levels were 200- to 300-fold higher after NM2CAB compared to N2CAB in rodents. Similarly, NM2CAB was slowly eliminated (t0.5 184 days) from plasma. CAB was detected in plasma for two years in monkeys.

In all species tested, NM2CAB and NCAB were widely distributed to all tissues with the exception of brain. The highest concentrations of both CAB and M2CAB were detected at the site of injection in muscle as well as in lymph nodes and spleen. After NM2CAB administration in mice, M2CAB concentrations were 10- to 100-times higher compared to CAB in lymph nodes and spleen and 50- to 100-times higher at the site of injection. Similar to plasma, CAB tissue levels were significantly higher after NM2CAB administration compared to NCAB in both mice and rats. Furthermore, the difference in tissue levels increased over time. By the end of the study, CAB levels in all tested tissues after NM2CAB treatment were up to 4,000-times higher compared to those after NCAB.

M2CAB levels in plasma and tissues are linked to levels of plasma CAB. However, plasma prodrug levels after NM2CAB injection were low, while tissue, including the site of injection, exhibited high prodrug and parent drug concentrations for the entire study period. In contrast, CAB tissue concentrations after NCAB administration rapidly declined to undetectable levels within five weeks. Therefore, the source of the high and sustained plasma CAB concentrations associated with NM2CAB administration is dependent on the rate of nanocrystal dissolution and prodrug release in tissue and at the injection sites and the hydrolysis rate of the ester chemical bond proved rapid after the surfactant coated nanocrystals was transformed into solution. Thus, it can be concluded that the sustained blood and tissue PK *in vivo* is due to the stability of NM2CAB and slow M2CAB dissolution at tissue and injection sites over time periods up to a year.

To confirm our PK findings, we repeated the male Balb/c mouse study in females of another mouse strain (NSG). In addition, this study was performed by three independent laboratories (the UNMC College of Pharmacy, the UNMC College of Medicine^15^, and Covance Laboratories). Finally, samples generated from the Covance Balb/c mouse study were quantified by Covance and by our laboratory using completely different LC-MS/MS methods. Results from all these studies were in agreement. CAB plasma concentrations were initially higher up to 6 weeks after NCAB administration followed by a faster decline compared to NM2CAB. CAB was detected for the entire period of the study after NM2CAB administration, while it was generally undetected or around the detection limit by 3 months after M2CAB administration. At all time points after 6 weeks where CAB was detected, its concentration was 100- to 1,000-fold higher after NM2CAB compared to NCAB administration. The t_0.5_ of CAB was extended from 3-8 days to 65-131 days after NM2CAB administration compared to NCAB. Finally, reproducible results were obtained for the study samples analyzed by two independent laboratories using different LC-MS/MS methods. Collectively, this provides conclusive evidence that the enhanced PK properties of NM2CAB over at least one year is reproducible in both sexes and in two strains of mice (Balb/c and NSG) by three different laboratories and is independent of the bioanalysis.

In summary, we demonstrate both rigor and reproducibility of our year-long CAB prodrug formulation. NM2CAB administration to two strains of mice, rats, and monkeys resulted in enhanced and sustained CAB plasma levels for at least one year. Plasma concentrations were up to, or greater than, 100-times those resulting from the equivalent NCAB. We also elucidated the mechanisms behind this enhanced PK profile from tissue distribution and *in vitro* data. NM2CAB accumulates and is retained in the injection site and in tissues, which serve as depots that slowly release the prodrug over months to years. *In vitro* data suggests that the observed enhanced tissue accumulation of NM2CAB is due to enhanced cellular uptake of the nanocrystals from the injection site, intracellular retention and nanocrystal stability in macrophages. In tissues, M2CAB undergoes slow dissociation which then is rapidly hydrolyzed into native CAB by both chemical and enzymatic hydrolysis. This was cross-validated by chemical and biological stability studies *in vitr*o using blood and tissue homogenates. Overall, the integration of mechanistic *in vitro* and *in vivo* data sets has elucidated the mechanisms behind the enhanced and sustained PK behavior of NM2CAB. These data sets support the premise that enhanced macrophage nanocrystal delivery and formation of intracellular tissue drug depots and slow prodrug dissolution rate contribute to the extended NM2CAB apparent drug half-life.

## Online Materials and Methods

### Chemicals

CAB was purchased from BOC sciences (Shirley, NY). Methanol (LC/MS Grade), acetonitrile (LC/MS Grade), water (LC/MS Grade), formic acid (FA, LC/MS Grade), ammonium formate (AF), bovine serum albumin (BSA), and phosphate buffered saline (PBS) were obtained from Fisher Scientific (Fair Lawn, NJ). Blood from mouse, rat, rabbit, monkey, dog and human was purchased from Innovative Research (Novi, MI). Dulbecco’s Modification of Eagle’s Medium (DMEM) was purchased from Corning Life Sciences (Tewksbury, MA).

### Nanoformulation Preparation and Characterization

Both CAB (NCAB) and M2CAB prodrug (NM2CAB) nanoformulations were prepared in a poloxamer 407 (P407) surfactant solution (10:1 ratio P407:drug) in endotoxin free water by high-pressure homogenization as previously described ^15^ in the Nebraska Nanomedicine Production Plant (NNPP) using good laboratory practices (GLP) protocols. The formulations were characterized for particle size, polydispersity index (PDI), and zeta potential by dynamic light scattering (DLS) using a Malvern Zetasizer Nano-ZSP (Worcestershire, UK). Drug loading was determined by ultra-performance liquid chromatography tandem mass spectrometry (LC-MS/MS) ^15^. Endotoxin concentrations were determined using a Charles River Endosafe nexgen-PTS system (Charles River, USA) and only formulations with endotoxin levels less than 5 EU/kg were used for animal studies.

### Storage Stability

Chemical stability was assessed by drug release measurements performed at room temperature. Two concentrations were studied including the original formulation after manufacturing (neat; 55.4 mg/ml for NM2CAB and 36.7 mg/ml for NCAB) and 20-fold (20X) diluted formulation (2.77 mg/ml for NM2CAB and 1.83 mg/ml for NCAB) in water. The neat and 20X dilutions were used for the rodent and *in vitro* studies, respectively. Nanoformulations were stored at room temperature and samples were collected on days 0, 4, 7, 14, 30, 60, 90, 252 and 365. For quantification of the released drug, aliquots were collected from the above samples and diluted in 4% BSA to adjust the drug concentration to 1 µg/ml, then samples were centrifuged at 10,000 g for 10 minutes to pellet and separate nanoparticles from the released/uncoated drug in the supernatant. Fifty µl aliquots were collected before and after centrifugation in 7 volumes of methanol (containing 0.1% FA and 2.5 mM AF). CAB from NCAB incubations and M2CAB from NM2CAB incubations were quantified by LC-MS/MS. Formulations were aliquoted in separate vials at day 0. The first were the actual incubations, where samples are taken at the various time points from one incubation to measure total drug concentration in the system before centrifugation as well as supernatant concentration after centrifugation in 4% BSA as described above. The second represented mass balance and contained 18 vials, enough for 2 vials to be extracted for total drug content at every time point. Stability was calculated as total amount of drug in the system at all time points: same at day 0. Release was calculated as amount of drug in supernatant: total amount of drug in the incubation (total before centrifugation) at each time point.

## Biological Stability

### Blood

To determine differences in rate of CAB and M2CAB stability in blood across species, 100 µl mouse, rat, rabbit, dog, monkey, and human blood was incubated with 1 µM native-CAB, native-M2CAB, NCAB, or NM2CAB at 37 °C. At different time points (0, 30 min, 2 h and 6 h), 0.9 ml of methanol (containing 0.1% FA and 2.5 mM AF) was added to each sample and vortexed for 3 min. The samples were then centrifuged at 16,000 g for 10 min after which 10 µl of the supernatant was mixed with 80% methanol (containing 0.1% FA and 2.5 mM AF) containing an internal standard (IS) and analyzed by LC-MS/MS. For the time zero start time, a 100 µl of ice-cold blood was spiked with M2CAB and immediately 0.9 ml of methanol (containing 0.1% FA and 2.5 mM AF) was added. Heat-inactivated plasma was incubated at the same conditions and used as negative controls to differentiate chemical vs. biological instability.

### Tissue

For metabolic stability in tissues, freshly collected rat liver, spleen, muscle, and lymph node tissues were homogenized in five-volumes of PBS (w/v). One hundred µl tissue homogenates were incubated with 1 µM M2CAB or NM2CAB at 37 °C. At different time points (0, 30 min, 2 h and 6 h), 0.9 ml of methanol (containing 0.1% FA and 2.5 mM AF) was added to each sample and vortexed for 3 min. Samples were centrifuged at 16,000 g for 10 min, 10 µl supernatant was mixed with 80% methanol (containing 0.1% FA and 2.5 mM AF) containing IS and analyzed by LC-MS/MS.

### Liver S9 Fraction

The metabolic stabilities of CAB and M2CAB were determined using mouse, rat, rabbit, dog, monkey, and human liver S9 fractions (XenoTech, LLC, Lenexa, KS, USA) following standard protocols ^28^. Briefly, CAB and M2CAB were incubated at 1 µM in a 100 µl mixture containing S9 fractions at 1 mg/ml protein concentration, NADPH (1 mM), saccharolactone (5 mM), uridine 5′-diphospho-glucuronic acid (UDPGA) (1 mM), and 3′-phosphoadenosin-5′-phosphosulphate (PAPS) (0.1 M) in PBS (100 mM, pH 7.4) at 37 °C. For each time point (0, 30, and 120 min), reactions were quenched by adding 0.9 ml of methanol (containing 0.1% FA and 2.5 mM AF), followed by centrifugation at 15,000 g for 10 min. Ten µl of the supernatants were then analyzed by LC-MS/MS. Heat-inactivated S9 fractions were incubated at the same conditions and used as negative controls. Hydroxycoumarin (7-HC) metabolites including 7-HC-glucuronide and 7-HC-sulfate, and the testosterone metabolite 6β-hydroxytestosterone were used as positive controls for phase I and phase II metabolism.

### Cellular Uptake, Retention and Drug Release

Human peripheral blood monocytes were obtained by leukapheresis from HIV-1,2 and hepatitis B seronegative donors and purified by centrifugal elutriation. Monocytes were cultured for the first seven days in macrophage colony stimulating factor enriched media (MCSF, 1000 U/ml) to facilitate cell vitality and differentiation into macrophages ^15, 29^. Monocyte-derived macrophages (MDM) were cultured in DMEM containing 4.5 g/L glucose, L-glutamine, and sodium pyruvate, and supplemented with 10% heat-inactivated human serum, 50 µg/ml gentamicin, and 10 µg/ml ciprofloxacin. Cells were maintained in clear flat-bottom 12-well plates at 37 °C in a 5% CO2 incubator at a density of 1.0 x 10^6^ cells per well. Half-culture media was replaced with fresh media every other day. After differentiation, MDM were utilized for CAB prodrug or drug nanoparticle uptake, retention and release. To quantify cellular uptake and retention kinetics, MDMs were incubated with 10 µM native-CAB, NCAB, native-M2CAB or NM2CAB. For uptake, MDM were collected at 8 h following treatment to measure intracellular drug and prodrug levels. For retention studies, after 8 h uptake, loaded MDMs were washed twice with PBS and fresh culture medium was added, and half-media was replaced every other day until the cells were harvested. At days 1, 5, 10, 15, 20, and 30, adherent MDM were washed twice with PBS, scraped into PBS, and counted using an Invitrogen Countess Automated Cell Counter (Carlsbad, CA). Cell suspension was then centrifuged at 900 g for 8 min at 4 °C. Cell pellets were sonicated in 200 µl of 80% methanol (containing 0.1% FA and 2.5 mM AF) to extract intracellular drug. The resultant lysates were centrifuged at 15,000 g for 10 min at 4 °C and supernatants were analyzed for CAB and M2CAB contents by LC-MS/MS. To quantify release kinetics, drug concentrations were measured in samples collected from the culture media from the same retention experiment using preloaded MDMs, at the same time when cells were harvested to quantify intracellular drug retention.

### Animal dosing and sampling

Eight-week-old, healthy male Balb/c mice and Sprague-Dawley (SD) rats were purchased from Charles River Laboratories (Wilmington, MA). Both mice and rats were housed in the University of Nebraska Medical Center (UNMC) laboratory animal facility accredited by the American Animal Association and Laboratory Animal Care (AAALAC). Mice were maintained on sterilized 7012 Teklad diet (Harlan, Madison, WI), and water was provided *ad libitum*. All procedures were approved by the Institutional Animal Care and Use Committee (IACUC) at the University of Nebraska Medical Center (UNMC) as set forth by the National Institutes of Health (NIH).

Two treatment groups of a single intramuscular injection (IM, caudal thigh muscle; 1.35 µl/g and 1.23 µl/g of body weight for mice and rats, respectively) of 45 mg CAB equivalents/kg of NCAB or NM2CAB were used for both mice and rats. For each treatment of either NCAB or NM2CAB, four groups of mice and rats (N=6) were dosed on day 0, and then at 1,3, 6, and 12 months each group was sacrificed and tissues including liver, lymph nodes, kidneys, spleen, lungs, brain, gut, muscle, and muscle from the site of injection were collected. In mice, inguinal, axillary, branchial, and cervical lymph nodes were collected and pooled, while in rats only cervical lymph nodes were collected. Following injection, blood samples were collected at 4h, days 1, 2, 4, 7, 14, 28, 42, and 56 and then monthly for 12 months into EDTA tubes (blood sample time points were evenly distributed between the 4 groups). Fifty µl of blood was immediately transferred into 950 µl methanol (containing 0.1% FA and 2.5 mM AF), vortexed, and stored at −80 °C until drug analysis. The remaining portion of the blood samples were centrifuged at 2,000 g for 5 minutes for plasma collection, which was stored at −80 °C until drug analysis. A one-year PK study of NM2CAB, performed in male rhesus monkeys (RM), was previously described ^15^. We report here PK assessment for samples collected for an additional year. Plasma samples were collected up to day 672 and CAB and M2CAB quantification were performed by LC-MS/MS.

### Covance PK studies

To provide added rigor and reproducibility to the data sets affirmation PK studies were performed by an independent contract laboratory. Male Balb/c and female NOD.Cg-*Prkdc^scid^ Il2rg^m1Wjl^*/SzJ (NSG) mouse studies were performed by Covance Laboratories (Greenfield, IN, USA). Briefly, 11-week-old male Balb/c and female NSG mice were purchased from Envigo RMS Inc. (Indianapolis, IN) and Charles River Laboratories (Wilmington, MA), respectively. Both mouse strains were housed at Covance Laboratories in an AAALAC-approved facility and the study was conducted using a Covance IACUC-approved animal protocol. Animal treatments, clinical observations, sample collections and sample analyses were conducted using standard operating procedures. Certified rodent diet #2014C (Envigo RMS, Inc.) was provided *ad libitum* to Balb/c mice and irradiated rodent diet #2920X (Envigo RMS, Inc.) was provided *ad libitum* to NSG mice. NCAB and NM2CAB formulations were prepared in the NNPP using GLP protocols. Formulation endotoxin levels were below 5 EU/kg. Formulations were packaged and shipped overnight at ambient temperature (18.1°C average) to Covance Laboratories in Greenfield, IN. Drug concentrations in the formulations were determined by UPLC-MS/MS at UNMC and by UPLC-UV/Vis at Covance. Personnel involved in drug administration, sample collection and sample analysis were blinded as to the treatments. Animals were divided into eight treatment groups that received NCAB or NM2CAB (N=6) as a single IM injection (1.3 µl/g, caudal thigh muscle) of 45 or 70 mg CAB equivalents/kg. Blood samples were collected via submandibular puncture into tubes containing lithium heparin from day 14 and every other week thereafter, up to 183 days. These studies mirrored previously reported studies performed by the Gendelman laboratory at UNMC in male Balb/c and female NSG mice ^15^. Plasma samples were stored at −80°C until analysis. Frozen plasma aliquots from each animal at each timepoint were shipped on dry ice by overnight shipping to UNMC for drug analysis. Samples received at UNMC were stored at −80°C until analysis.

### Sample Preparation and LC-MS/MS Analyses

Blood and tissue sample preparation and analysis were performed as previously described [17]. For blood samples, 50 µl blood collected in 950 µl of methanol was vortexed, centrifuged at 16,000 g for 10 min at 4 °C, and 50 µl supernatant was aspirated, mixed with 50 µl IS [20 ng/ml d3-dolutegravir (d3- DTG) and 40 ng/ml stearoylated darunavir (SDRV)] in 70% methanol (containing 0.1% FA and 2.5 mM AF). d3-DTG was used as the IS (final concentration = 10 ng/ml) for the quantification of CAB, and SDRV (final concentration = 20 ng/ml) was used as the IS for quantification of M2CAB. Ten µl of sample was analyzed by LC-MS/MS for CAB and M2CAB. For tissue sample preparation, 3-150 mg of each tissue sample were homogenized in 4-35 volumes (depending on the tissue) of 80% methanol (containing 0.1% FA and 2.5 mM AF) using a TissueLyzer II (Qiagen, Valencia, CA, USA). Two hundred and ninety µl of methanol (containing 0.1% FA and 2.5 mM AF) and 10 µl 80% methanol was added to 100 µl of tissue homogenates. Samples were vortexed for 3 min, and centrifuged at 16,000 g for 10 min. Then, 50 µl supernatant was aspirated and mixed with 50 µl IS in 80% methanol (containing 0.1% FA and 2.5 mM AF). Ten µl sample was injected on LC-MS/MS for CAB and M2CAB analysis. Calibration curves in the range of 0.05-500 ng/ml for CAB and MCAB were prepared the same way in blank blood and tissues.

For LC-MS/MS quantification of CAB and M2CAB, a Waters ACQUITY UPLC system (Waters, Milford, MA, USA) connected to a Waters Xevo TQ-XS mass spectrometer with an electrospray ionization source was used, as described previously with some modifications ^15^. For CAB analysis, chromatographic separation was achieved on an ACQUITY UPLC BEH Shield RP18 column (2.1×100 mm, 1.7 µm; Waters) using 7-min gradient of mobile phase A (7.5 mM AF, pH 3) and mobile phase B (100% acetonitrile) at a flow rate of 0.25 ml/min. The initial mobile phase composition was 37% B for the first 4.0 min, increased to 95% B over 0.25 min, and held constant for 1.25 min. Mobile phase B was then reset to 37% over 0.5 min and the column was equilibrated for 1 min before the next injection. M2CAB chromatographic separation was achieved on ACQUITY UPLC BEH Shield RP18 column (2.1×30 mm, 1.7µm; Waters) using 8-min gradient of mobile phase A (7.5 mM AF, pH 3) and mobile phase B (100% methanol), at a flow rate of 0.28 ml/min. The initial mobile phase composition was 85% B for the first 5 min, increased to 95% B over 0.25 min, and held constant for 1.5 min. Mobile phase B was then reset to 85% over 0.25 min and the column was equilibrated for 1 min before the next injection. CAB, M2CAB, d3-DTG, and SDRV were detected at a cone voltage of 2 V, 4 V, 2 V and 70 V, respectively, and a collision energy of 22 V, 20 V, 16 V and 16 V, respectively, in the positive ionization mode. Multiple reaction monitoring (MRM) transitions used for CAB, M2CAB, d3-DTG, and SDRV were 406.21 > 127.08, 672.47 > 406.16, 423.27 > 277.21, and 814.70 > 658.61, respectively. Spectra were analyzed and quantified by MassLynx software version 4.1. All calculations were made using analyte to IS peak area ratios.

### Blood Drug Measurements

Mean blood drug concentrations were calculated per treatment group and PK parameters were derived using non-compartmental analysis of blood concentration vs. time profiles, using Phoenix WinNonlin software (version 8.2). Peak plasma concentration (C_max_), time to reach C_max_ (T_max_), elimination rate constant (λ^Z^), half-life from elimination phase (t_0.5_), area under the plasma concentration versus time curve (AUC), mean resident time (MRT), clearance (CL/F), and apparent volume of distribution (Vz/F) were calculated. Tissue concentrations were calculated and expressed as ng/g tissues.

## Supporting information

Gautam et al Supplemental Figures

## Supplementary Material

(See attached document)

## Acknowledgments

We thank the University of Nebraska Medical Center (UNMC) Elutriation and Cell Separation Core (Myhanh Che and Na Ly) for providing human monocytes. We are thankful to Benjamin G. Lamberty, Brenda M. Morsey and Dr. Howard Fox for the extension of previously published data sets performed with NM2CAB in rhesus macaques (in Nat Mater. 2020 Aug;19(8):910-920). This work was supported, in part, by the University of Nebraska Foundation, which includes donations from the Carol Swarts, M..D. Emerging Neuroscience Research Laboratory, the Margaret R. Larson Professorship, the Frances and Louie Blumkin and the Harriet Singer Research Donations. We thank Dr. Bradley Britigan, Dean of the College of Medicine at UNMC, for providing funds to support the Covance studies. The authors thank Sarah Favara, Emily Archer and the Covance Laboratory staff that assisted in conducting and analyzing the data sets for rigor and reproducibility. We thank Dr. Jennifer Larsen, the Vice Chancellor for Research for UNMC for Core research support enabling continuance of this work. The research also received support from National Institutes of Health grants PO1 DA028555, R01 NS36126, PO1 MH64570, P30 MH062261, R01 AG043540 and 2R01 NS034239. We also thank the INBRE grant support from 2P20GM103427 for infrastructure research support.

## Author contributions

N.G. Designed the study, executed the laboratory and animal PK experiments, performed the data analysis and interpretation, co-wrote the manuscript; J.M. Assisted in study design, data analysis and interpretation, co-designed and coordinated the Covance study and plasma sample analyses, edited the manuscript and assisted in the preparation of the figures; D.K. Performed the animal PK experiments, responsible for the data acquisition and analysis; A.B. Assisted in animal study design and co-executed the animal PK experiments; Q.P., W.L. Assisted with the animal PK sample preparation and analysis and performed LC-MS/MS assays; T.K. Assisted in human monocyte cell culture experiment and data analysis. N.S. Assisted in animal PK sampling; B.L.D.S. Performed LC-MS/MS analyses of all samples from the Covance study and assisted in data analysis from the study; B.S. Synthesized the prodrug and prepared and characterized the nanoformulations for both the College of Pharmacy and Covance studies and assisted with LC-MS/MS tests of Covance mouse plasma samples; A.S. Prepared and characterized all formulations for the College of Pharmacy and Covance studies; B.E. assisted in study design, created nanoformulation, and edited the manuscript; H.E.G. Co-developed the study design, co-wrote the manuscript, developed the infrastructure responsible for Nebraska Nanomedicine Production Plant for the nanoformulations used in study, developed the Covance research design and provided funding support for the studies; Y.A. Was responsible for the study design and data interpretation and co-wrote the manuscript.

