## Supplementary material for "Lipophilic Nanocrystal Prodrug-Release Defines the Extended Pharmacokinetic Profiles of a Year-Long Cabotegravir": Gautam et al Supplemental Figures

**Supplementary Figure Legends:**

**Supplementary Figure 1. Blood CAB and M2CAB concentrations.** Blood concentration vs. time profiles for CAB and M2CAB over one year after a single IM injection (45 mg CAB equivalent/kg) of NCAB or NM2CAB in (A) mice and (B) rats. Data are expressed as mean ± SEM (N=6 animals).

**Supplementary Figure 2. Bioanalytical confirmation of mouse plasma CAB concentrations.** Male Balb/cJ or female NSG mice were given a single intramuscular injection of 70 mg CAB equivalents/kg of NCAB or NM2CAB. Plasma was collected every other week for 6 months at which time the study was terminated. CAB concentrations were quantitated at UNMC by LC-MS/MS. Data are expressed as mean ± SEM (N=3-6 animals per treatment group).

**Supplementary Figure 3. M2CAB pH stability.** Chemical stability profile of M2CAB at pH 1.0 (0.1M HCl), pH 7.4 (PBS), pH 8.3 (heat inactivated plasma) and pH 11.0 (0.1M NaOH) at 1µM drug concentration. (A) Per cent M2CAB remaining vs. time profile for M2CAB; (B) Per cent CAB formation vs. time profile from M2CAB.

**Supplementary Figure 4. Plasma stability of M2CAB and NM2CAB.** Metabolic stability profiles of M2CAB and NM2CAB three species (mouse, rat, and rabbit), and metabolic stability profiles of CAB and NCAB in six species (mouse, rat, rabbit, monkey, dog, and human) and heat-inactivated plasma at 1 µM concentration in blood. Data shown as

M2CAB disappearance over time for (A) M2CAB and NM2CAB; CAB formation from (B) M2CAB and NM2CAB as well as from (C) CAB, and (D) NCAB.

**Supplementary Figure 5. Liver S9 stability of M2CAB.** Metabolic stability profile of M2CAB at 1μM concentration in liver S9 fraction from six species (mouse, rat, rabbit, monkey, dog, and human) and heat-inactivated liver S9 fraction. (A) Per cent M2CAB remaining vs. time profile for M2CAB; (B) Per cent CAB formation vs. time profile from M2CAB.

**Supplementary Figure 6. Tissue stability of M2CAB and NM2CAB.** Metabolic stability profiles of M2CAB, and NM2CAB at 1 μM concentration in rat tissues (liver, spleen, muscle, lymph node, and heat-inactivated liver). Data shown as M2CAB disappearance over time for (A) M2CAB and (C) NM2CAB and metabolite (CAB) formation from (B) M2CAB and (D) NM2CAB.

**Supplementary Figure 7. NM2CAB formulation stability.** Stability was tested on three separate NM2CAB formulations sitting on the benchtop at room temperature after manufacture for the Covance 45 mg CAB equivalents/kg (61.8 mg/ml), Covance 70 mg CAB equivalents/kg (89.9 mg/ml) and UNMC College of Pharmacy (COP; 55.4 mg/ml) PK studies. Formulation stability was measured by particle hydrodynamic diameter (size), polydispersity index (PDI), and zeta potential as determined by dynamic light scattering (DLS). All results are shown as the mean of three replicates.

Supplementary Figure 1

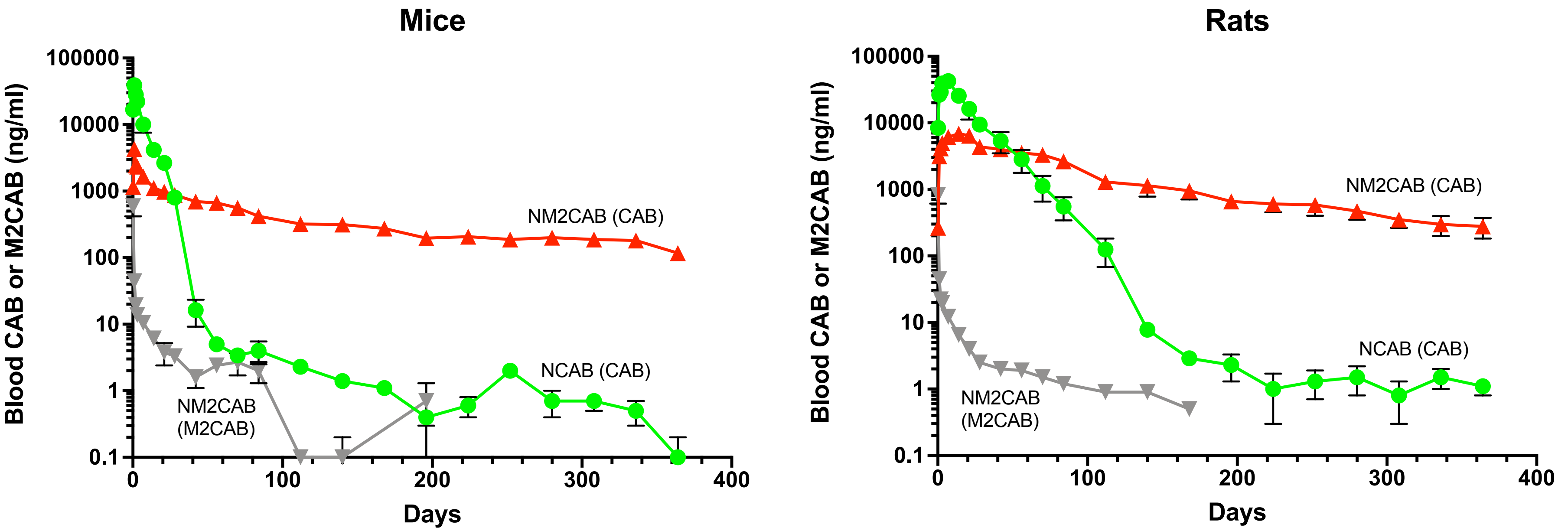

Supplementary Figure 2

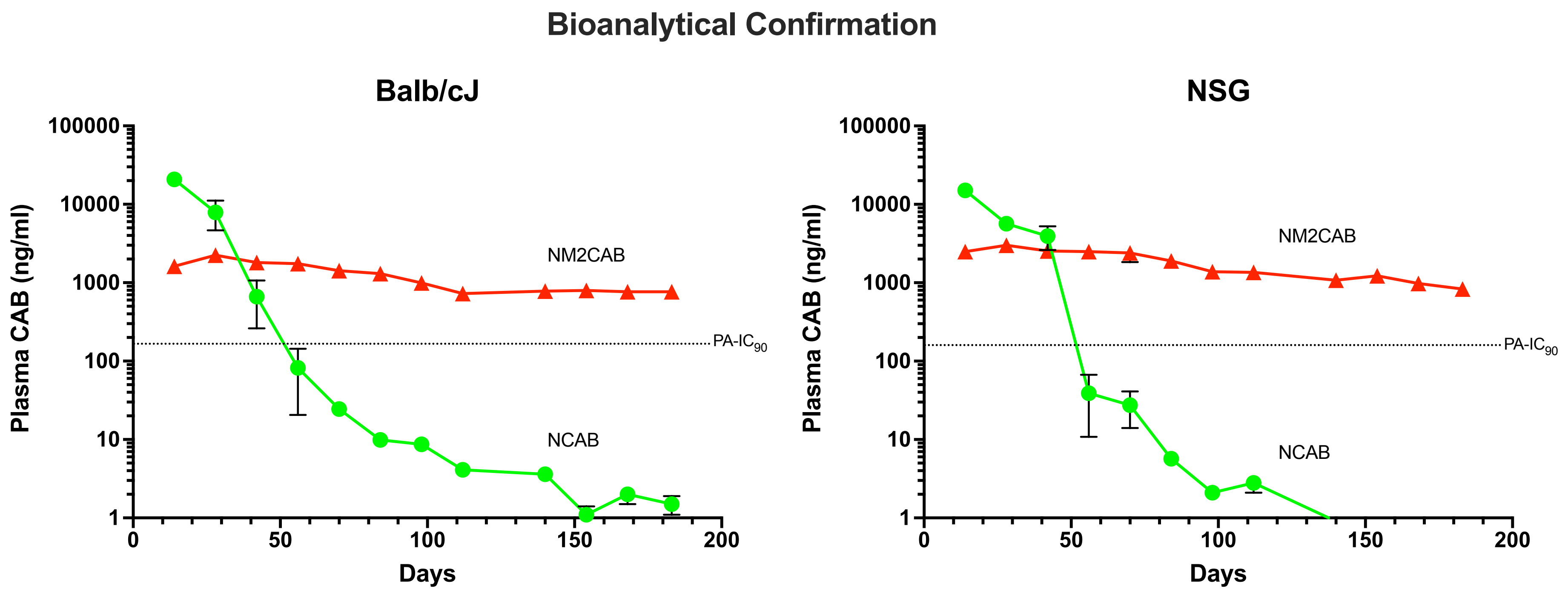

Supplementary Figure 3

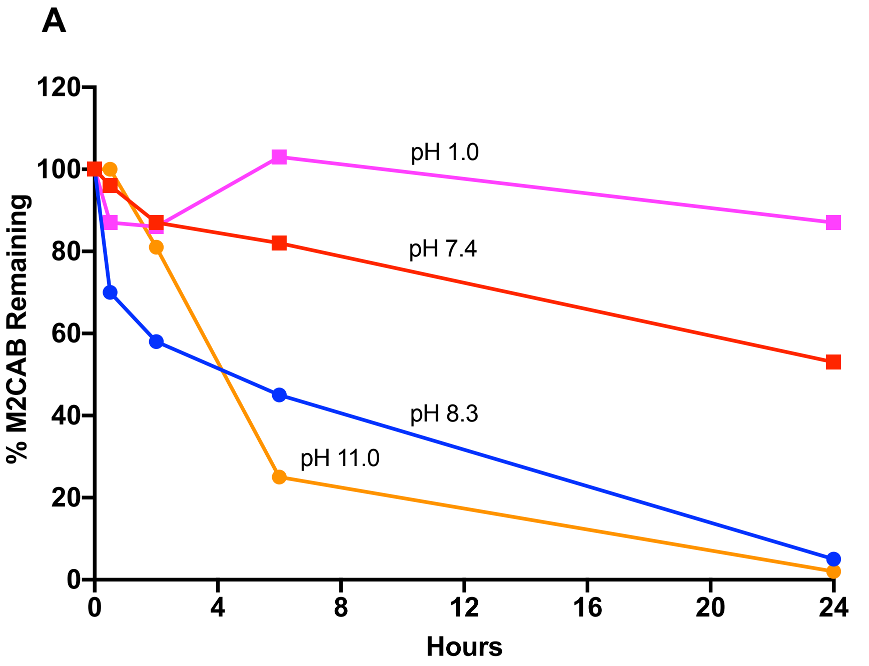


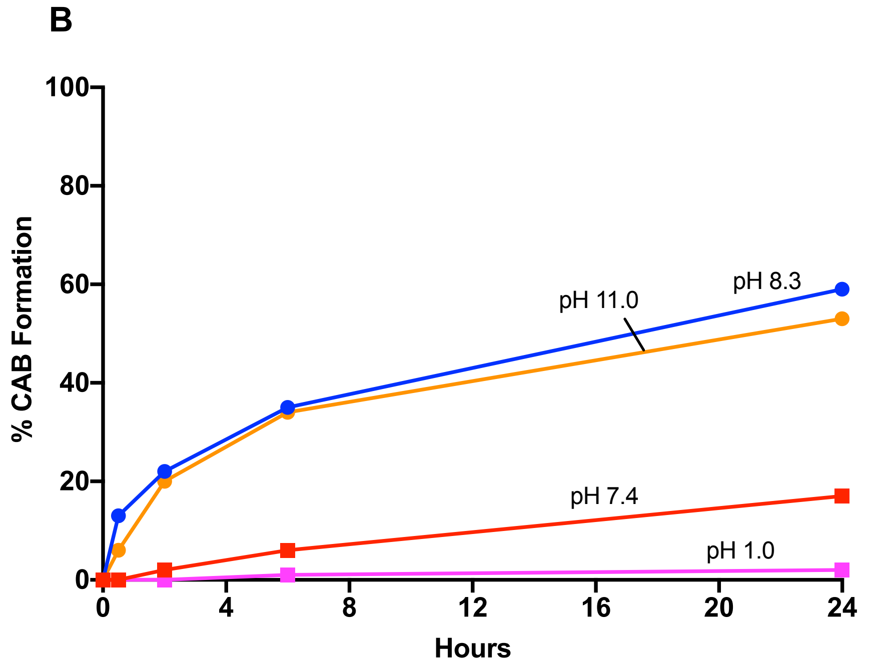

### Supplementary Figure 4

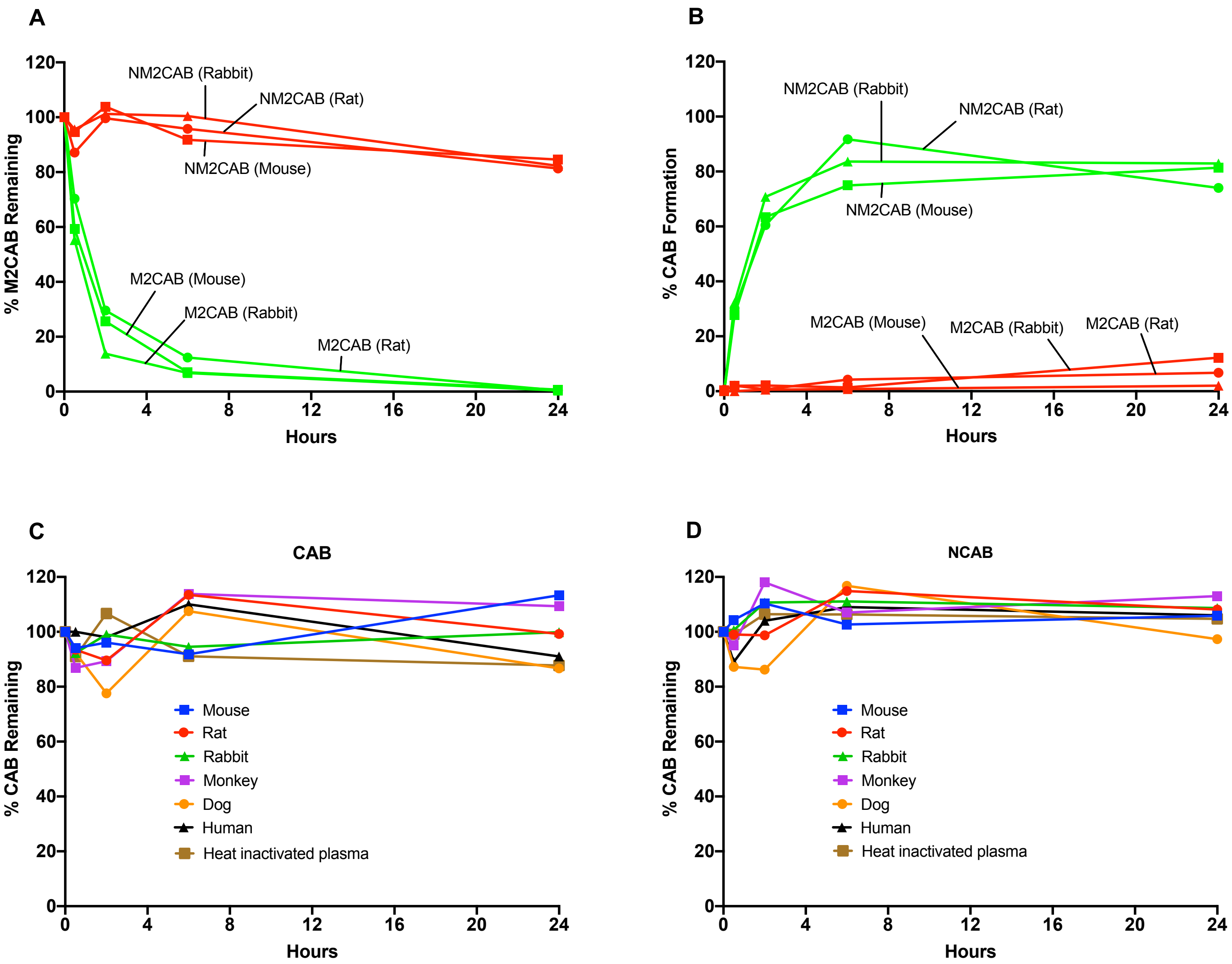

Supplementary Figure 5

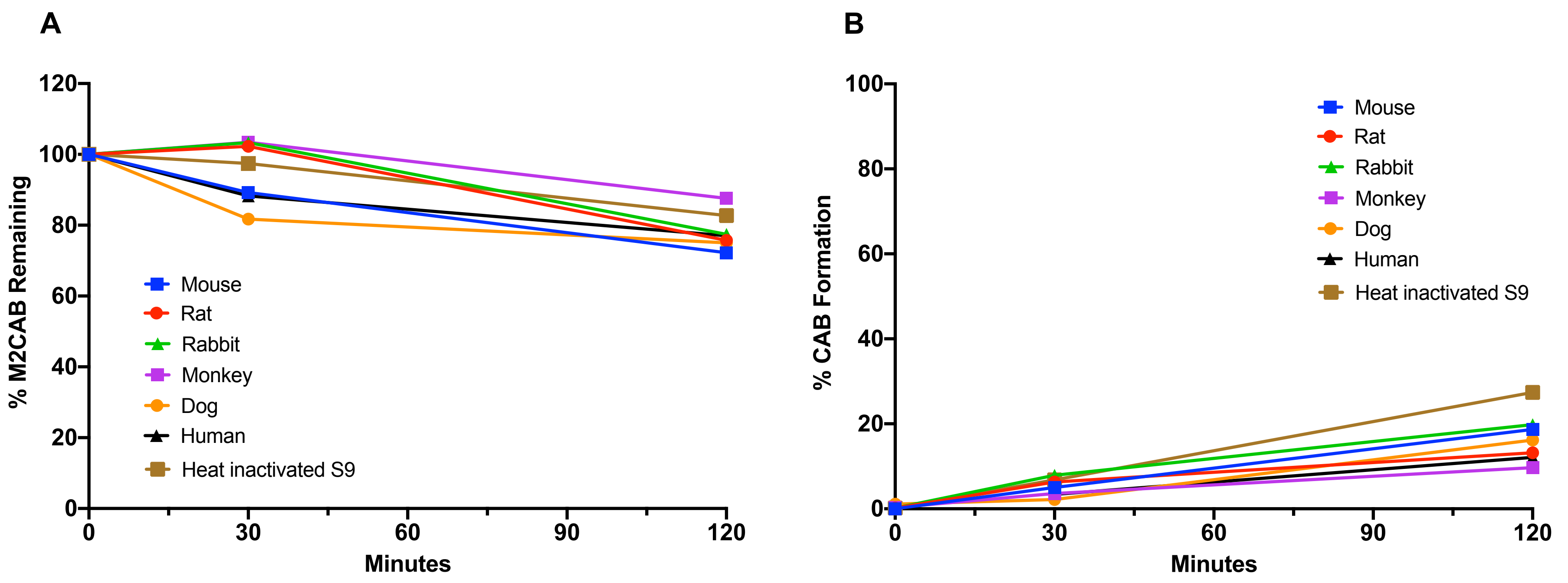

Supplementary Figure 6

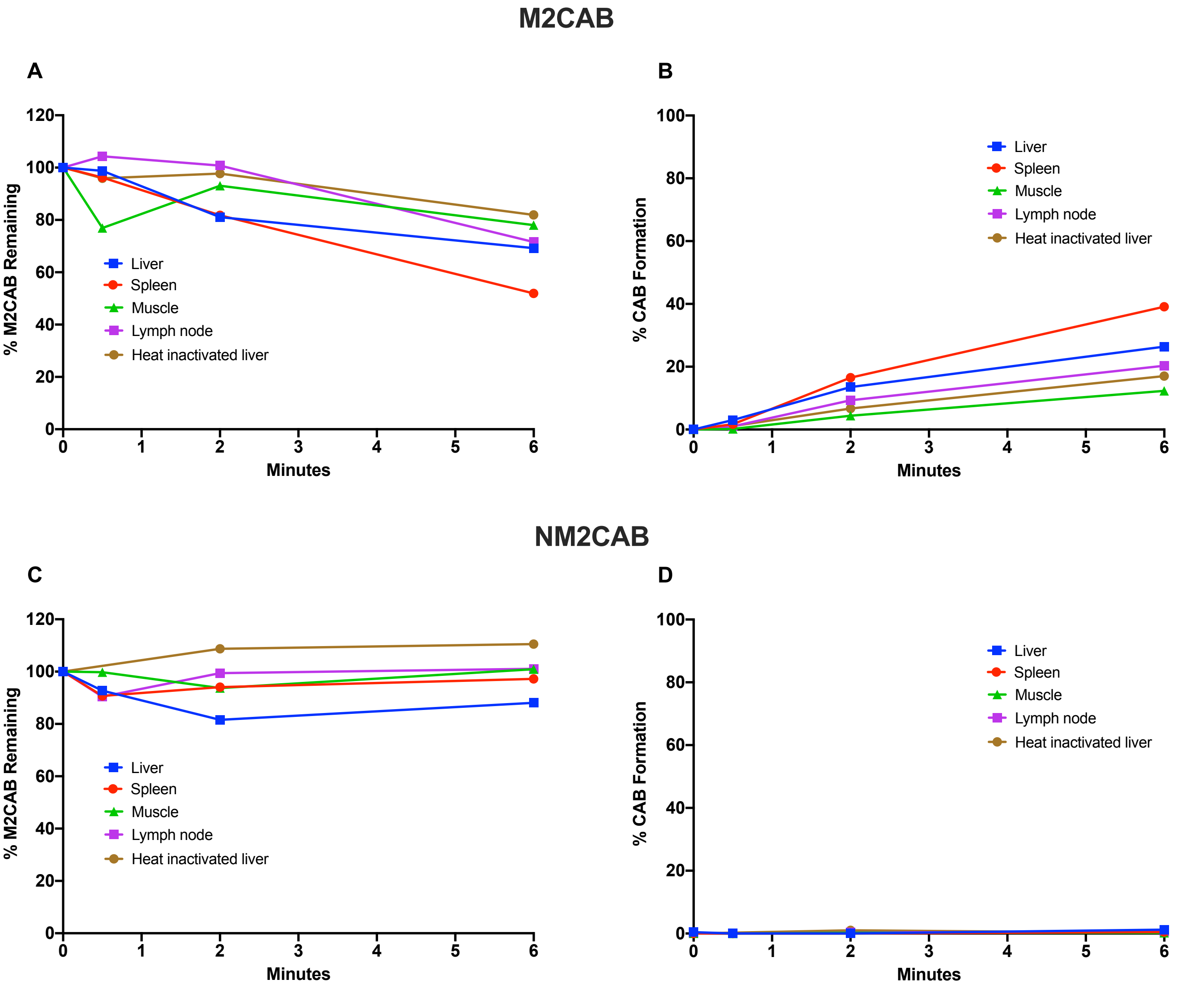

Supplementary Figure 7

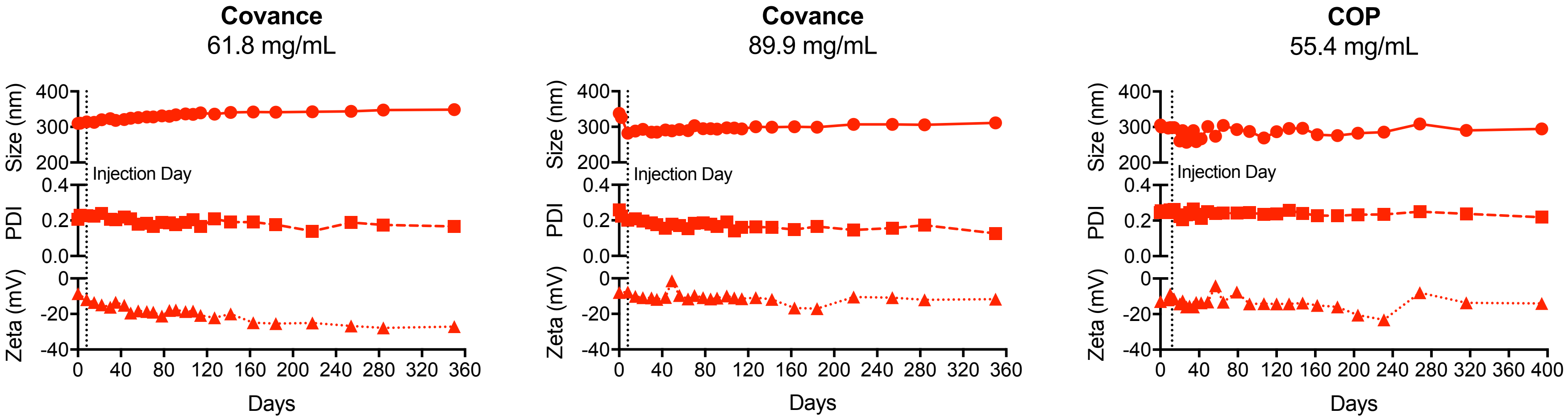
